# Neurodegeneration emerges at a cellular tipping point between aggregate accumulation and removal

**DOI:** 10.1101/2025.09.08.674880

**Authors:** Matthew W. Cotton, Shriram Venkatesan, Joseph S. Beckwith, Dorothea Böken, Catherine K. Xu, Emre Fertan, Jonathan C. Breiter, Lexie E. Berkowicz, Laura Sancho Salazar, Alex Von Schulze, Ewa A. Andrzejewska, Emma E. Brock, Hannah L. Han, Matthias M. Schneider, Danny D. Sahtoe, David Baker, James B. Rowe, Alain Goriely, William A. McEwan, Tuomas P.J. Knowles, Steven F. Lee, Randal Halfmann, David Klenerman, Georg Meisl

## Abstract

Protein aggregates are a pathological hallmark across neurodegenerative diseases. Yet, the disconnect between molecular-level aggregation and the emergence of disease severely limits mechanistic understanding of neurodegeneration. Here, we bridge this disconnect by showing that a cellular tipping point emerges as a universal feature across diseases from the competition between aggregate accumulation and removal. We map the resulting cellular phase transition with our high-throughput live-cell assay, measuring the tipping point that separates healthy cells from those with large aggregate loads. Using super-resolution imaging of brain tissue from Alzheimer’s and Parkinson’s disease, we quantify how the balance of accumulation and removal is shifted in disease. We validate our framework by predicting how designed aggregation inhibitors shift the tipping point to restore cellular homeostasis. Our results provide a mechanistic framework connecting molecular-level aggregation to disease, paving the way for a quantitative, unified understanding of neurodegeneration and enabling predictions of therapeutic efficacy.

---

The vast majority of neurodegenerative diseases are characterised by the conformational conversion of normally soluble proteins into pathological aggregates (*1, 2*), a simple biophysical process that occurs even in purified proteins, without the need for any cellular factors. Despite this perceived simplicity, the processes that control the emergence and progression of disease remain poorly understood. A prevailing hypothesis is that aggregates act in a prion-like fashion, where a single misfolded aggregate seeds a cascade of aggregation. This hypothesis has been proposed as an explanatory framework for the spreading of pathology throughout the brain, but provides limited insight for guiding therapeutic development for the most common neurodegenerative diseases, such as Alzheimer’s disease (AD) and Parkinson’s disease (PD).

Crucially, this seeded self-replication alone cannot explain two central observations: First, ageing, and the associated progressive decline of proteostasis and aggregate removal capacity, is a dominant risk factor (*3–6*). Second, aggregates also occur in healthy brain regions and individuals (*7–9*). Recent single-molecule detection and super-resolution imaging studies reveal substantial populations of tau aggregates, which form in AD (*10, 11*), and disease-associated *α*-synuclein aggregates, which form in PD (*12, 13*), in samples from healthy adult human brains (see Fig. 1a, b). These observations imply that organisms can tolerate, or even limit aggregation for extended periods, and point to additional control layers, such as active cellular mechanisms (chaperones, proteasome, autophagy–lysosomal pathways) that continually counteract the ongoing aggregate accumulation. An alternative hypothesis then emerges, that, in addition to the ability of aggregates to self-replicate (*14*), an imbalance of aggregate production and removal is the crucial step in the emergence of disease (*15*). The opposing effects of aggregation and removal may lead to non-linear dynamics and a tipping point beyond which uncontrolled proliferation of aggregates, *runaway aggregation*, is inevitable (*16*) (see *Auxiliary Supplementary Materials Cotton et al. 2025*).

**Figure 1.**
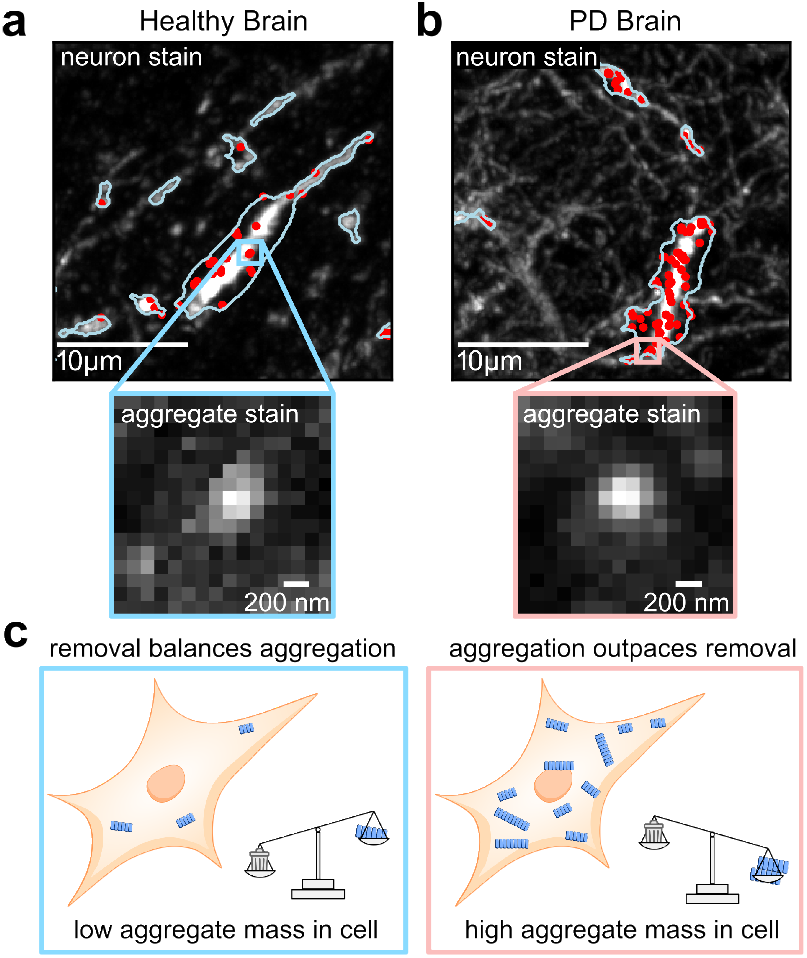
A binary cells state with aggregates in healthy controls. Direct imaging of human brain tissue reveals aggregates in cells in both (**a**) healthy control brains and (**b**) brains from individuals with Parkinson’s disease. The figure shows exemplary images of cells containing aggregates (red points) where neuron cell boundaries (blue outline) are imaged using a Microtubule-Associated Protein 2 (MAP2) stain and aggregates are imaged using a phosphorated U-synuclein (pS129) stain, insets. (**c**) A schematic of the tipping point between aggregation and removal. See SI for imaging details.

In this work, we test these hypotheses and confirm that the balance between protein aggregate production and removal leads to a cellular tipping point between health and disease as a universal feature. Cells exist in one of two distinct states, a healthy state characterised by low aggregate levels, or a pathological state with an abundance of protein aggregates (Fig. 1c). This explains the predominantly binary cell state seen in histological brain images and cell culture studies (*12,17–19*). A critical transition point separates the two states and, by considering active removal mechanisms in vivo, we are able to use a mathematical model to predict and experimentally verify how this tipping point changes depending on physiology or therapeutic interventions.

## Quantifying the reduced rate of aggregate removal in AD and PD

We first investigate the molecular signatures of a changed balance between aggregate production and removal in disease. Impaired aggregate removal has been proposed as a key driver in the onset and progression of many neurodegenerative diseases (*3–6*). Genome-wide association studies (GWAS) have linked increased genetic risk to components of the aggregate removal machinery, including the ubiquitin–proteasome system and microglial function (*20*). Additionally, protein quality control declines with age (*6, 21, 22*), reducing the cell’s ability to remove aggregates.

High resolution, single-aggregate imaging techniques now enable the detailed measurement of many thousands of sub-µm-sized aggregates from human samples, producing size distributions that contain information about the relative rate at which aggregates are removed compared to the rate at which they are formed, see Fig. 2a,b,c. This single aggregate resolution is crucial as the shape of the larger clumps of aggregates detected by conventional histopathology is predominantly governed by their surroundings, such as the cell shape, and therefore contains little information on the molecular processes that govern their formation.

**Figure 2.**
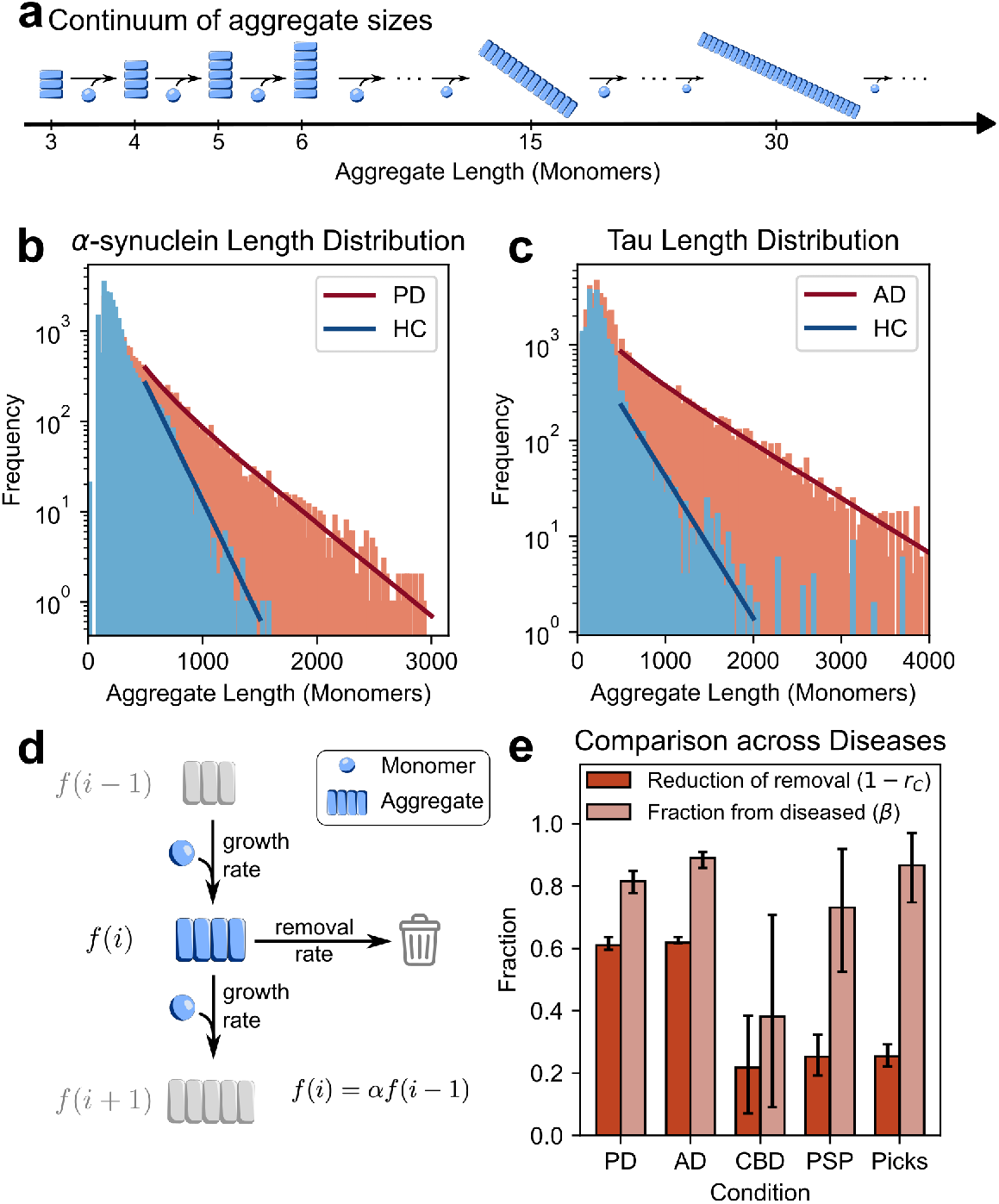
Aggregate length distributions reveal reduction in removal mechanisms in disease. **(a)** Super-resolution imaging reports a continuum of aggregate sizes at different lengths. **(b)** Length distribution of small *α*-synuclein aggregates from post-mortem measurements in healthy brains (n=3) and brains with PD, stage 4-5 (n=3) (frontal cortex) (*25*). **(c)** Tau aggregate length distribution from super resolution SiMPull (see SI for TIRF micrographs) measurements of brain homogenate from healthy (n=5) and AD cases (n=5) (frontal cortex) (*11*). The distributions in both (b) and (c) are well described by our model fits, solid lines, predicting significantly reduced removal in disease. **(d)** Schematic for a minimal elongation-removal model of aggregate size. **(e)** Summary of the reduction in relative removal rates, 1 – *r*_*C*_, and the proportion of aggregates from pathological cells, *β*, for PD, AD and a number of tauopathies (for CBD n=5, PSP n=5, Picks n=4). The 2 fits for each dataset have a total of 3 free parameters: relative removal rates in healthy and diseased cells and proportion of aggregates from diseased cells. n is the number of individuals.

To extract this mechanistic information, we develop a mathematical model that accurately describes the observed aggregate size distribution and quantitatively links its shape to the balance between aggregate growth and removal. Specifically, when aggregates are long enough for their growth rates to no longer be size-dependent (*23, 24*), the aggregate size distribution in a cell is expected to follow a geometric decay *p*_*i*_ ∝ *α*^*i*^, where an aggregate of size *i* occurs with frequency *p*_*i*_, as illustrated in Fig. 2d. The decay factor, *α*, is determined by the ratio of the rates of aggregate growth and removal (see SI for details of model and fits). Intuitively, if aggregate removal is impaired, aggregates will persist for longer and grow larger, yielding a slower decay in the length distribution. Thus, changes in *α* provide a direct readout of the balance between aggregate removal and growth. Indeed, using high-resolution imaging of protein aggregates in postmortem human brain tissue, we see exactly this geometric decay in the aggregate length distribution (see straight lines on the logarithmic plots in Fig. 2b,c arising from the distributions in the healthy brain samples). In diseased brains however, the situation is more complex, with some cells still in their stable healthy state but a significant proportion of cells in the pathological state. The measured aggregate distribution thus reflects a mixture of two populations, resulting in curved lines on the logarithmic length distribution plots (Fig. 2b,c, PD and AD distributions). We quantify this difference by defining *r*_*C*_, the ratio of the removal rate in pathological cells compared to healthy cells (see SI for derivation), and *β*, the fraction of aggregates from pathological cells. We show this as the reduction in the relative removal rate in disease, 1 – *r*_*C*_, in Fig. 2e, so that more severe disease corresponds to higher values of both 1 – *r*_*C*_ and of *β*. We apply this analysis to size distribution measurements of *α*-synuclein aggregates in brains from individuals with PD and to tau aggregates from brains of individuals with AD and other tauopathies (see SI for details) (*10, 12*). The fits are shown in Fig. 2b,c, results are summarised in Fig. 2e.

As predicted, we find a higher proportion of aggregates at larger sizes in diseased brains. Remarkably, in cases where the distribution has significant contributions from both healthy cells and those with pathological protein aggregation, see PD in Fig. 2b, there is a clear evidence of the predicted curvature in the length distributions, matched accurately by our model.

This analysis allows us, from patient data, to quantify both the degree to which the molecular processes are altered in disease, and the proportion of the aggregates that came from cells in this altered pathological state. In PD, we find that 82% [78% – 85%] of the *α*-synuclein aggregates are from pathological cells and in those cells, the relative removal is reduced by 61% [60% – 64%] compared to healthy cells (mean with 68% confidence intervals from Bayesian inference, see SI for details). By contrast, for tau in AD we find that the removal rate is reduced by 62% [62%–64%], but 89% [86%–91%] of the aggregates are from pathological cells. The other tauopathies, Corticobasal Degeneration (CBD), Pick’s disease (Picks), and Progressive Supranuclear Palsy (PSP), show a less pronounced, albeit still significant, reduction in the rate of removal.

### Minimal model yields a cellular tipping point as a universal feature

Having demonstrated the ability of an aggregate production and removal model to extract mechanistic information from human data, we now consider a minimal model to determine the key features of such a system. The balance between aggregate production and removal can be quantitatively explained by a mechanistic model with two key ingredients: (1) an aggregate formation process, whose rate depends on the concentration of available monomeric protein (*23, 26*) and (2) an aggregate removal process, with a maximum rate of removal. We remain agnostic to the specific aggregation mechanisms in order to establish the general system behaviour that emerges robustly across realisations. A maximum rate is the only stipulation on the removal process and is justified by the fact that the cellular machinery responsible for aggregate removal has limited capacity, due to the energy demands of the process, finite concentrations of enzymes and other limitations (*27, 28*).

The system is described by the total protein concentration, *m*_tot_, and the fraction of the protein in the cell that is in an aggregated state, *µ*. Aggregation occurs at some rate *A* and removal at some rate *R*. Both can in principle depend on the monomer and aggregate concentrations and so the evolution of the aggregate mass in a cell becomes

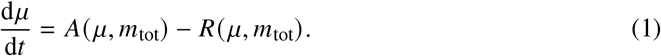

Whether the mass of aggregates in the cell increases or decreases depends on whether the aggregate production or removal rate is larger, i.e. the relative magnitudes of *A* and *R*. This results in distinct steady-state behaviours (the fixed points of the system) determined by the relative balance of the two processes.

When removal is dominant, the model predicts that the system will converge to the low level steady state aggregate mass, where aggregates are cleared from the cell at the same rate at which they are formed, *R* = *A*, and thus the system remains in balance. We identify this as the state of healthy cells: they have a non-zero aggregate mass that is regulated by cellular removal mechanisms. For typical parameter values, this healthy steady-state aggregate concentration is extremely low. By contrast, when the balance between protein aggregation and removal is disrupted, the system enters runaway aggregation. We identify these cells as ‘pathological’, although multiple additional downstream mechanisms may be involved in generating pathology.

The exact molecular mechanisms that drive aggregation and removal remain elusive and likely differ between diseases and specific aggregating proteins. However, the emergence of two stable states, and how to transition between them, is a robust phenomenon observed across diverse models (see *Auxiliary Supplementary Materials Cotton et al. 2025*). As a representative example, we consider a specific realisation of the dynamics of equation (1) where the aggregation includes a primary nucleation, secondary nucleation and an elongation process (with rate constants *k*_n_, *k*_2_ and *k*_+_ respectively). This gives

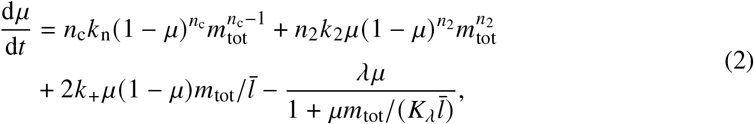

where *ī* is the average aggregate size and λ is the rate constant of removal at low aggregate concentration. This assumes a chaperone-mediated removal of aggregates, with maximum rate *V*_max_ = λī*K*_λ_ and Michaelis constant *K*_λ_. The primary and secondary nucleation processes have reaction orders *n*_c_ and *n*_2_ respectively (see SI for full derivation).

A phase-plane plot of this system (Fig. 3a) illustrates how monomer and aggregate concentrations determine the cell’s fate. In this plot, arrows indicate the direction of change in aggregate mass—upward for increasing, downward for decreasing—while the solid lines denote steady states where aggregate production is balanced by removal. When a cell’s aggregate mass and monomer concentration fall within the shaded pale blue region, the system evolves towards the healthy, stable fixed-point line. However, when the state lies within the red region, the production of aggregates outpaces the cell’s removal capacity, driving the system into a regime of uncontrolled, pathological aggregation. States within this red region ultimately converge toward the high-aggregate–mass ‘pathological’ state. The dashed black line is the transition boundary between these two regions. Mathematically, this is the line of unstable, repulsive fixed points, terminating at the two bifurcation points.

**Figure 3.**
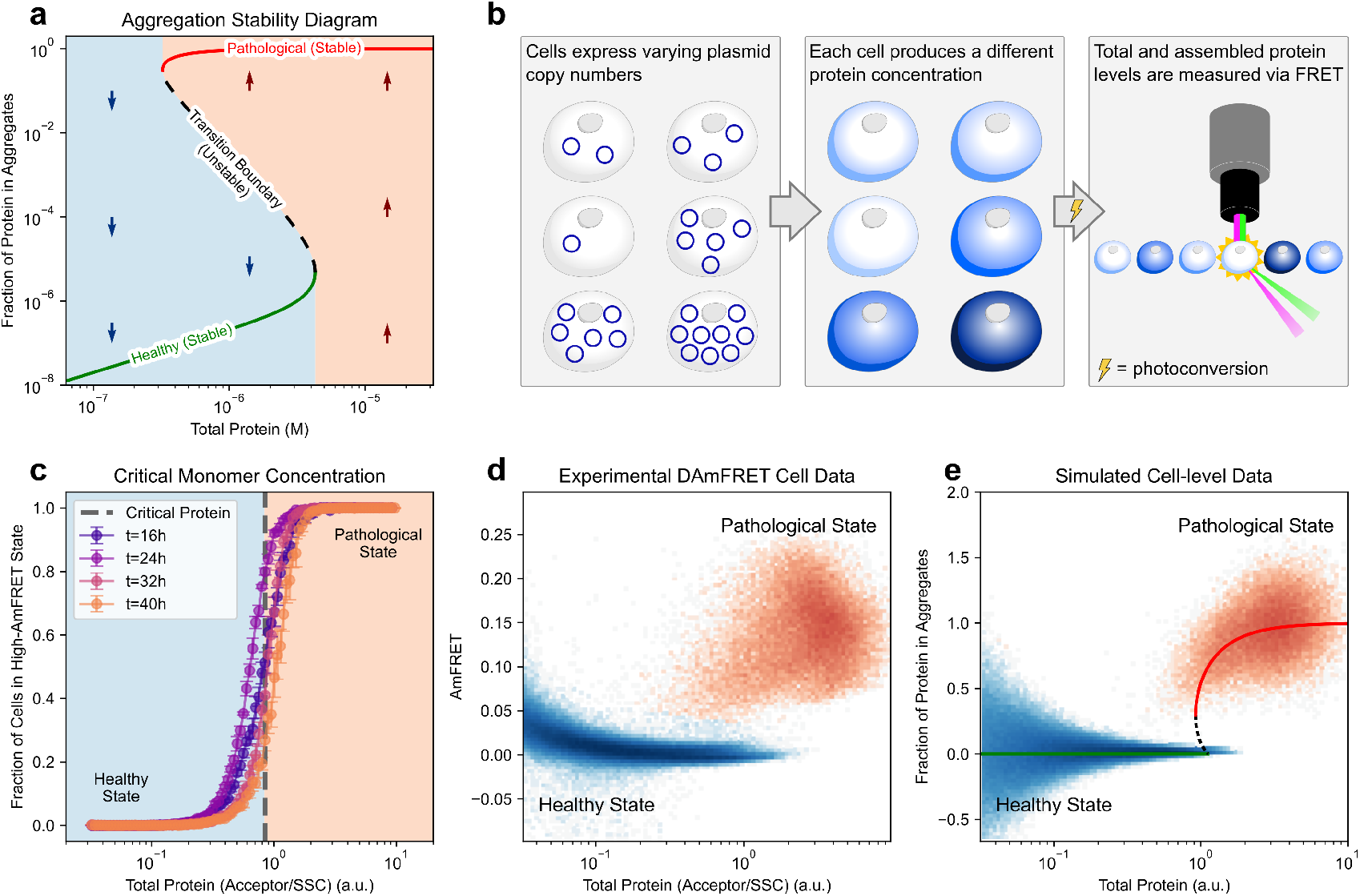
DAmFRET reveals bistability in cells. (**a**) Phase plane analysis of a minimal model, equation 2, showing whether the system will have a net production of aggregates (upwards arrows), or net removal of aggregates (downwards arrows). The two stable branches (solid lines) and an unstable transition branch (dashed line) are shown. The pale blue region shows the set of cell states that will evolve to the healthy state (the basin of attraction). System parameters given in SI. (**b**) Schematic of the DAmFRET technique used to measure total protein and aggregate concentrations in a cell population. (**c**) The proportion of cells that contain mostly aggregated protein shows a sudden transition around a critical monomer concentration that is constant in time. (n=3 at each timepoint, error bars are the standard deviation, for individual repeats see SI) (**d**) DAmFRET data of cells expressing a model amyloid show two clear populations – high and low AmFRET – and shows good agreement with the simulated data in (e). (**e**) Example simulated distribution of cell states based on the fixed points of the model. The solid/dashed lines indicate stable/unstable fixed points for an average cell. Differences in the shape of the healthy state population are due to how noise is modelled in simulations. For further details and simulation parameters see SI.

The shape of the transition boundary highlights an important prediction: the tipping point can be crossed by either an increase in the protein concentration (shift along the x-axis) or by an increase in the aggregate concentration (shift along the y-axis). The former leads to an imbalance by increasing the rate of aggregate formation, as the formation rate depends on the monomer concentration (*23, 26*). The latter leads to an imbalance by increasing the demands on the removal system beyond its abilities to cope, overwhelming the aggregate removal mechanisms. Above the critical monomer concentration, the rate of aggregate formation always exceeds the maximum aggregate removal capacity, and the system inevitably progresses toward pathological aggregation, even if the aggregate fraction is reduced (*16*). The shape of this phase plot also has an important implication for therapeutic interventions. Once cells have transitioned to the pathological state, it is significantly more difficult to move them back to their healthy state than it is to prevent the transition in the first place. This hysteresis demonstrates that minimal interventions may be an effective form of prevention, even if they are not sufficient to treat established disease.

### A high-throughput cell assay reveals the phase plane of aggregation

In order to validate this model, we explored the phase plane of aggregation experimentally. Distributed amphifluoric FRET (DAmFRET) (*29*) is a powerful method to simultaneously measure monomer and aggregate concentrations in individual cells. Each experiment contains a population of ~10^5^ cells harbouring random numbers of plasmids such that the expressed protein’s concentration varies by a factor of over 100 between cells. Fig. 3 shows the DAmFRET data for a model amyloid (HET-s 218-289) expressed in yeast cells and reveals that the cells form two distinct populations: a low aggregate state at low protein expression and a high aggregate state at high protein expression.

While normally used to investigate nucleation behaviour (*29*), we rule out that the tipping point in our specific cell model is driven by a stochastic waiting time by performing time course measurements of the same cell system. If the tipping point were dominated by slow primary nucleation kinetics or governed by a stochastic waiting time then even at low monomer concentrations, cells would eventually transition to an aggregated state, albeit over long timescales. However, we find that the distribution is unchanged over time, implying that each of the two states is stable, which is consistent with active processes working maintain the low aggregate state, see Fig. 3c.

These DAmFRET data offer an exceptional approach to directly measure the aggregation phase plot. As predicted, we find that two populations emerge, one at low protein concentrations with low amounts of aggregates, and one at high protein concentrations with a high amount of aggregates, with a sudden transition at the critical monomer concentration separating the two, Fig. 3d. Using equation (1), we can simulate the cellular aggregate concentrations in a large population of cells and we recover extremely good agreement with the experimentally measured DAmFRET data as shown in Fig. 3e.

This empirical dataset not only validates our model, it also highlights the importance of protein concentration in determining the state of the cell. The transition to an aggregating state by increasing the protein concentration is particularly important in many animal models, where overexpression of the disease-relevant proteins is used to trigger aggregation in a system where it would normally not occur (*30*). In humans, increased expression levels are associated with increased prevalence or earlier onset (*31*), such as increased A*β* expression in adults with Down’s syndrome causing AD (*32–34*) or triplication of the gene locus which produces *α*-synuclein causing familial PD (*35, 36*). Our model now allows quantitative linking of these changes in expression levels to the cell state.

### Limited tolerance to seeds points to maximum aggregate removal capacity

Even cells that are otherwise in a healthy state can be driven into pathology if a sufficiently large mass of aggregates is introduced, overwhelming the clearance machinery. This scenario corresponds to a vertical crossing of the stability line in Fig. 3a.

This transition predicts that the sudden addition of aggregate mass into a cell, if large enough, can cause a cell to cross the tipping point. The key difference to more common descriptions of seeding, which simply assume that the cell is in a kinetically trapped state until a single seed is introduced, is the presence of a critical mass of seeds. Adding an amount larger than this critical mass will increase the aggregation rate enough to overwhelm the removal mechanisms and trigger runaway aggregation (see Fig. 4a). However, if a lower than critical mass of additional aggregates is introduced to a cell, the system will not cross the tipping point and the removal mechanisms will still be able to return the cell to a healthy state. These two scenarios describe super- and sub-critical seeding, respectively, and are illustrated schematically in Fig. 4a and on the phase plane in Fig. 4b. This plot also illustrates that the closer the system is to its critical point, the fewer aggregates are required for super-critical seeding. If the stability line is moved, for example by reducing removal through stressing the system, a reduction in the number of seeds required for super-critical seeding can occur, see Fig. 4c. In contrast, nucleation-limited models do not exhibit sub-critical seeding behaviour, as a single aggregate is sufficient to induce seeding. Classical prion diseases (e.g. Creutzfeldt-Jakob disease) are examples of such a nucleation limited-system where the onset of disease can be initiated by the introduction of a single pathogenic aggregate. Similarly, introduction of other highly infectious agents, such as viruses, do not show sub-critical seeding, Fig. 4d.

**Figure 4.**
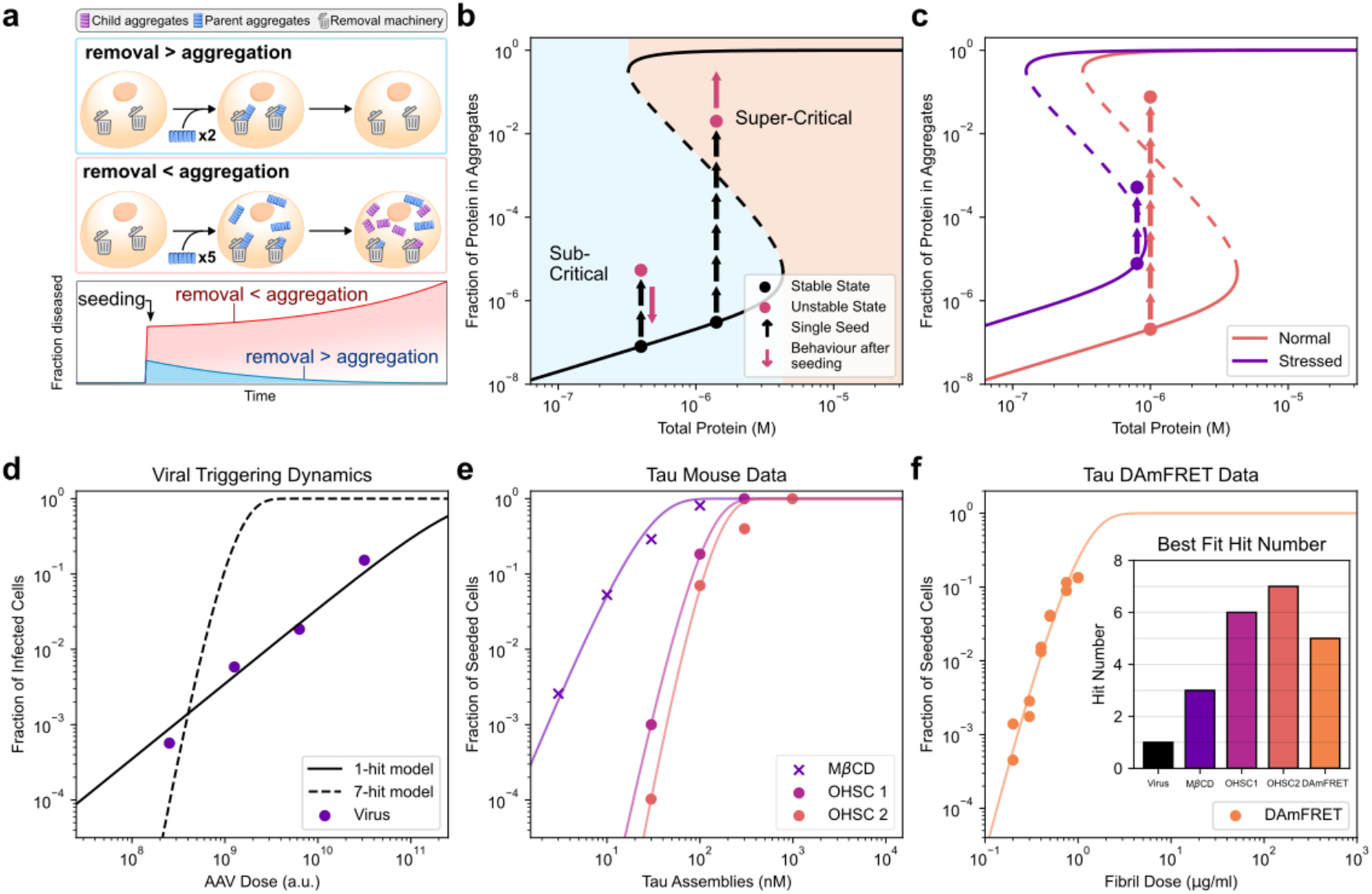
Effect of removal mechanisms on seeding. (**a**) Schematic showing the effect of seeding in systems with and without removal. (b,c) The phase plane illustrates how a cell can recover from a sub-critical seeding event, but how a super-critical seeding event can cross the critical boundary to trigger runaway aggregation, (**b**), and demonstrates how a shift in the phase plot changes the number of seeds required, (**c**). (**d**) Dose-response curve of Adeno-associated virus (AAV) infection of an OHSC follows one-hit dynamics. (e,f) The dose response curve of recombinant tau fibrils onto OHSCs (**e**) and tau reporter HEK cells (n=2 unique cell populations and approximately 10^4^ – 10^5^ cells per data point) (**f**). Inset summarizes the best fit values, except for AAV infection all systems show *h >* 1 hit dynamics, consistent with super-critical seeding. Raw data in (d) and (e) is from (*37, 38*) with slices from N= 6 mice per condition for OHSC1 and M*β*CD and N= 3 mice per condition for Viral and OHSC2.

Experimentally, measuring the scaling of the probability of seeding with seed concentration is a robust approach to determine whether seeding can be triggered by a single aggregate. When *h* seeds are needed to trigger runaway aggregation, the probability of seeding approximately follows an *h*^th^ order reaction *P*(seed) 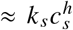 where *c*_*s*_ is the concentration of seeds and *k*_*s*_ is the rate of seeding (detailed derivation see SI). Plotting the fraction of pathological cells as a function of the seed concentration on a double logarithmic plot thus allows for easy and robust extraction of the number of seeds required to cause critical seeding in a cell: the slope corresponds to the number of seeds *h*, the position of the line is determined by the rate *k*_*s*_.

Fig. 4d,e,f shows such seeding dose response measurements for a number of different experimental systems. Panel d shows infection by a virus, a classical single-hit model requiring only a single virus to trigger infection (*37*). This is contrasted by the data in panels e and f, where assemblies of recombinant tau protein are added to organotypic hippocampal slice cultures (OHSCs) derived from P301S mice (panel e) or to a tau biosensor HEK293T cell line (panel f). The slopes of these curves all clearly exceed 1, for both the OHSC (*h* = 6–7) and the biosensor cells (*h* = 5), consistent with multiple seeds being required to trigger aggregation. Additionally, the OHSC data demonstrate that fundamental changes to cellular state can disrupt aggregate removal, move the transition line and alter the cell’s seeding threshold, *h*. This is illustrated in Fig. 4e, where seeding titrations onto OHSCs were performed in the presence and absence of the cholesterol-extracting agent methyl-beta-cyclodextrin (M*β*CD) (*38*). This treatment depletes membrane cholesterol and increases cytosolic entry of applied tau assemblies potentially bypassing some of the removal mechanisms that are active upon tau aggregates. Treatment with M*β*CD lowers the number of aggregates required for critical seeding from *h* = 6 to *h* = 3, consistent with predictions from our model for a scenario in which the system is moved closer to its critical point by the introduction of a cell stress and reduction of the removal capacity, Fig. 4c. Following a similar mechanism, stressing of the removal machinery may also explain the increased aggregation that has been observed in the presence of an inflammatory challenge (*39, 40*).

While we here modelled this overwhelming of removal processes by simply assuming a limited capacity of the removal pathways, there is also evidence for other feedback processes, where the presence of aggregates directly slows removal (*41*). Regardless of the specifics of this feedback, whether by limited capacity or by damage to the cellular machinery by existing aggregates, the fundamental behaviour of the system remains the same. A limited capacity for removal also naturally explains the emergence of co-aggregates and co-pathologies: aggregates of one protein slow generic removal processes, thus moving other proteins closer to their respective tipping points.

## Effect of aggregation inhibitors and therapeutic outlook

In addition to explaining monomer dependence, seeding response, and the changes in aggregate size distribution in disease, the model developed here can be used to predict the effect of therapeutic intervention and helps categorize therapies into (1) those that increase aggregate removal, and (2) those that reduce aggregation rates. For example, antibody based therapeutics, such as the anti-amyloid antibodies licensed for AD (*42–44*), aim to increase the removal rate of aggregates, whereas aggregation inhibitors (Fig. 5a) or monomer-reducing therapies, such as epigenetic editors (*45*) and the antisense oligonucleotides currently in clinical trials (*46*), aim to decrease the aggregate production rate. In both cases, therapeutic efficacy is determined by the intervention’s ability to restore the system to a stable state, i.e. shifting it to the pale blue region of the phase plot in Fig. 3a, either by moving the position of the system, or by moving the tipping point, Fig. 5b.

**Figure 5.**
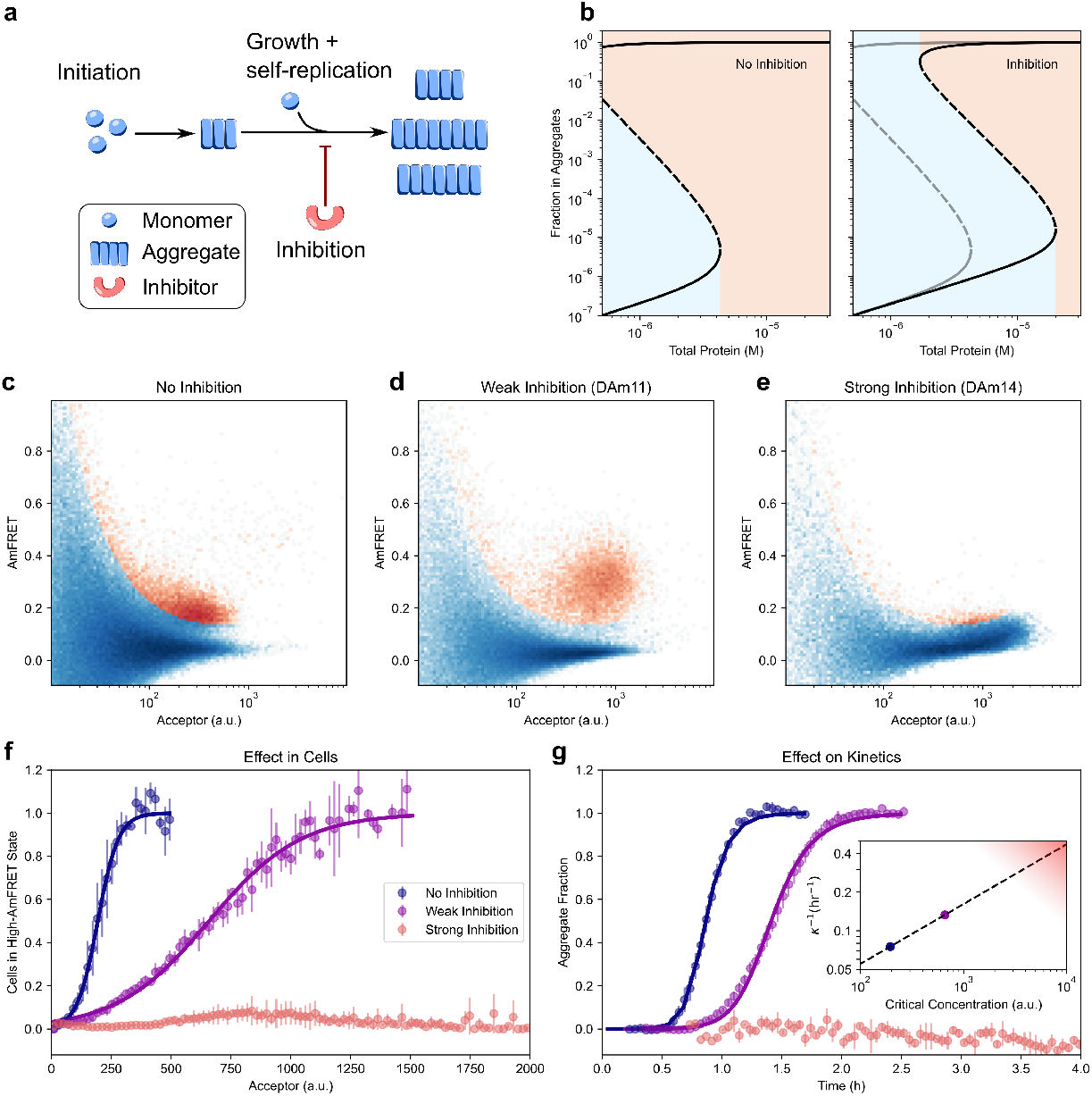
In vitro kinetics predict the shift in the cellular tipping point. (**a**) Schematic of the mechanism of binder inhibition. (**b**) Inhibition of self-replication by a factor of 100 moves the transition boundary and increases the healthy stable region. See SI for all parameters. (c-e) DAmFRET plots showing the distribution of cells expressing A*β* without inhibition (**c**), or with coexpressed weak (**d**) or strong (**e**) inhibitor. The 2D histogram is coloured to highlight the boundary between the two populations that is used in the subsequent analysis (see SI). (**f**) Shows the amount of cells that have transitioned to an aggregated state at each monomer concentration for different levels of inhibition (n=3). (**g**) Kinetic data from in vitro aggregation can be used to determine the strength of the inhibitors (data for strong inhibition and no inhibition from Sahtoe et al. (*47*), n=3). Inset: effective rate from in vitro aggregation kinetics and critical protein concentration from DAmFRET experiments correlate across the two systems. Straight line fits to no and weak inhibition on a double logarithmic plot show a reaction order of 0.47 with respect to monomer concentrations in the cells. Strong inhibition by DAm14 prevented aggregation, allowing for only a lower bound for the quantities, as denoted by the shaded area, but is consistent with the expected trend.

To demonstrate experimentally the effect of aggregation inhibition on the transition boundary, we again utilise DAmFRET measurements. Along with amyloid *β*1-42 (A*β*), we now co-express a protein-based inhibitor from the same plasmid. This means the inhibitor is expressed together with the protein directly in the cell, and factors such as uptake of inhibitor do not need to be considered. These protein inhibitors were designed against A*β*, and previously demonstrated their ability to inhibit A*β* aggregation in vitro (*47*). We here compare the effect of a weak and a strong inhibitor, referred to as DAm11 and DAm14, respectively. The results of the DAmFRET measurements of cells expressing either no inhibitor, a weak inhibitor or a strong inhibitor are shown in Fig. 5c,d,e. As the strength of the inhibitor increases, the population of pathological cells (orange) is only observed at higher monomer concentrations, with no clear pathological population being visible with the strong inhibitor. This effect is more clearly visible when the fraction of cells in the pathological state is shown as a function of A*β* concentration in Fig. 5g.

In order to test the predictive power of our model, we used the aggregation kinetic measurements, outlined in Sahtoe et al. (*47*), which are shown in Fig. 5f. In that work we found that the designed proteins inhibit aggregate formation by interfering with growth and / or self-replication of existing fibrils, effectively altering the rate of aggregation, *κ*.

In our cellular model, the position of the critical point is also determined by *^*. If the inhibitor acts in the same way within cells, a double logarithmic plot of the aggregation rate extracted from in vitro aggregation studies against the critical monomer concentration determined by DAmFRET should give a straight line. The slope of this line is determined by the reaction order of the aggregation process with respect to monomer in the cells (for details see SI). Indeed, a clear correlation between the two quantities is observed as shown in the inset of Fig. 5g, giving a reaction order of 0.47. Although no exact reaction order for the cellular system is known, reaction orders at comparable monomer concentrations in vitro agree with this value, ranging between 0.5 and 1.5 depending on the exact buffer conditions (*48–50*).

## Conclusion

In summary, we present a unifying mechanistic framework that explains the onset and progression of neurodegenerative diseases as a kinetic phase transition. This cellular tipping point, when protein aggregation and removal are no longer in balance, emerges as a universal feature. Beyond matching data of length distributions, seeding and the effectiveness of aggregation inhibitors, our framework naturally provides a mechanistic foundation for a range of observations. Disease-associated mutations of aggregating proteins, increased expression levels, and the introduction of infectious material shift cells towards disease by increasing the rate of aggregate formation (*51, 52*). By contrast, factors such as chronic inflammation, co-pathologies and ageing can shift the tipping point by reducing the effectiveness of removal mechanisms. This explains the often late age of onset and common involvement of inflammation observed in disease (*53*). Similarly, therapeutic approaches can be classified as acting on either aggregate formation or aggregate removal. Strategies for reducing aggregate formation include antisense oligonucleotides to lower monomer expression levels (*46, 54*) and specific inhibitors of aggregation (*47, 55*). By contrast, anti-amyloid antibodies such as Aducanumab and Lecanemab, the first disease-modifying drugs approved for Alzheimer’s, aim to increase aggregate removal (*42, 56*). Moreover, the s-shaped stability curves predict that systems that are already in the runaway aggregation state require a much more significant intervention to return to stability, providing a causal mechanism as to why prevention may be more achievable than a cure.

Conceptually, our work unifies many key questions about neurodegenerative disease, by reframing them into a simple consideration of how the tipping point is affected. By enabling molecular level analysis and prediction, it paves the way for quantitative understanding of the drivers of disease and ultimately the design of more targeted therapies.

## Acknowledgments

We thank Bradley Hyman and Jorin Riexinger for insightful feedback.

## Funding

This work was supported by a UK Dementia Research Institute pilot award (G.M., M.W.C.); The Newman Foundation (M.M.S. and T.P.J.K.); Additional support was provided by the National Institute for Health Research Cambridge Biomedical Research Centre (NIHR203312: the views expressed are those of the authors and not necessarily those of the NIHR or the Department of Health and Social Care) and the Medical Research Council (MC UU 00030/14; MR/T033371/1); the National Institute of General Medical Sciences of the National Institutes of Health (Award Number R01GM130927, to R.H.) and the Stowers Institute for Medical Research. L.E.B. is supported by the Madison and Lila Self Graduate Fellowship at the University of Kansas. The funders had no role in study design, data collection and analysis, or manuscript preparation. The content is solely the responsibility of the authors and does not necessarily represent the official views of the funders.

## Author contributions

M.W.C., A.G. and G.M. developed the mathematical theory and analysis. S.V., J.S.B., D.Bö., E.F., J.C.B., L.E.B., L.S.S., A.V.S., E.A.A., E.E.B., H.L.H., M.M.S. and D.D.S. performed experiments and analyzed results. D.Ba., W.A.M., T.P.J.K., S.F.L., R.H., and D.K. conceived the experiments. G.M. conceived the project. M.W.C. and G.M. led writing the manuscript, all authors contributed to revising and editing the manuscript.

## Competing interests

G.M. is a consultant for WaveBreak Therapeutics. W.A.M. is a founder and consultant for Trimtech Therapeutics.

## Data and materials availability

Original data underlying this manuscript can be accessed from the Stowers Original Data Repository at https://www.stowers.org/research/publications/libpb-2570.

## Notes

### Competing Interest Statement

G.M. is a consultant for WaveBreak Therapeutics. W.M. is a founder and consultant for Trimtech Therapeutics.

### Summary of Updates

Updated abstract. Updated Figure 2 with additional dataset and updated the author list accordingly.

## References and Notes

1. F. Chiti, C. M. Dobson, Protein Misfolding, Functional Amyloid, and Human Disease. Annual Review of Biochemistry 75 (1), 333–366 (2006), eprint: https://doi.org/10.1146/annurev.biochem.75.101304.123901, doi:10.1146/annurev.biochem.75.101304.123901, https://doi.org/10.1146/annurev.biochem.75.101304.123901.

2. F. Chiti, C. M. Dobson, Protein Misfolding, Amyloid Formation, and Human Disease: A Summary of Progress Over the Last Decade. Annual Review of Biochemistry 86 (1), 27–68 (2017), eprint: https://doi.org/10.1146/annurev-biochem061516-045115, doi:10.1146/annurev-biochem-061516-045115, https://doi.org/10.1146/annurev-biochem-061516-045115.

3. J. Labbadia, R. I. Morimoto, The biology of proteostasis in aging and disease. Annual Review of Biochemistry 84 (1), 435–464 (2015), tex.eprint: https://doi.org/10.1146/annurevbiochem-060614-033955, doi:10.1146/annurev-biochem-060614-033955, https://doi.org/10.1146/annurev-biochem-060614-033955.

4. R. I. Morimoto, Cell-Nonautonomous Regulation of Proteostasis in Aging and Disease. Cold Spring Harbor Perspectives in Biology 12 (4), a034074 (2020), publisher: Cold Spring Harbor Laboratory, doi:10.1101/cshperspect.a034074, http://doi.org/10.1101/cshperspect.a034074.

5. Y. Aman, et al., Autophagy in healthy aging and disease. Nature Aging 1 (8), 634–650 (2021), publisher: Nature Publishing Group, doi:10.1038/s43587-021-00098-4, https://doi.org/10.1038/s43587-021-00098-4.

6. J. Hardy, V. Escott-Price, The genetics of neurodegenerative diseases is the genetics of agerelated damage clearance failure. Molecular Psychiatry 30 (6), 2748–2753 (2025), publisher: Nature Publishing Group, doi:10.1038/s41380-025-02911-7, https://doi.org/10.1038/s41380-025-02911-7.

7. S. L. DeVos, et al., Synaptic tau seeding precedes tau pathology in human alzheimer’s disease brain. Frontiers in Neuroscience 12, 267 (2018), doi:10.3389/fnins.2018.00267, https://doi.org/10.3389/fnins.2018.00267.

8. C. M. Moloney, V. J. Lowe, M. E. Murray, Visualization of neurofibrillary tangle maturity in Alzheimer’s disease: A clinicopathologic perspective for biomarker research. Alzheimer’s & Dementia 17 (9), 1554–1574 (2021), doi:10.1002/alz.12321, https://doi.org/10.1002/alz.12321.

9. M. S. LaCroix, et al., Tau seeding without tauopathy. Journal of Biological Chemistry 300 (1), 105545 (2024), doi:10.1016/j.jbc.2023.105545, https://doi.org/10.1016/j.jbc.2023.105545.

10. D. Böken, et al., Single-Molecule Characterization and Super-Resolution Imaging of Alzheimer’s Disease-Relevant Tau Aggregates in Human Samples. Angewandte Chemie International Edition 63 (21), e202317756 (2024), doi:10.1002/anie.202317756, https://doi.org/10.1002/anie.202317756.

11. D. Böken, et al., Small tau aggregates exhibit disease-specific molecular profiles across tauopathies (2025), doi:10.1101/2025.06.10.658934, http://doi.org/10.1101/2025.06.10.658934.

12. R. Andrews, et al., Large-scale visualisation of α-synuclein oligomers in Parkinson’s disease brain tissue. bioRxiv : the preprint server for biology (2024), publisher: Cold Spring Harbor Laboratory tex.elocation-id: 2024.02.17.580698 tex.eprint: https://www.biorxiv.org/content/early/2024/02/19/2024.02.17.580698.full.pdf,doi:10.1101/2024.02.17.580698, doi:10.1101/2024.02.17.580698,.

13. B. Fu, et al., RASP: Optimal single puncta detection in complex cellular backgrounds. The Journal of Physical Chemistry B 128 (15), 3585–3597 (2024), tex.eprint: https://doi.org/10.1021/acs.jpcb.4c00174, doi:10.1021/acs.jpcb.4c00174, https://doi.org/10.1021/acs.jpcb.4c00174.

14. G. Meisl, et al., Uncovering the universality of self-replication in protein aggregation and its link to disease. Science Advances 8 (32), eabn6831 (2022), doi:10.1126/sciadv.abn6831, https://doi.org/10.1126/sciadv.abn6831.

15. G. Meisl, The thermodynamics of neurodegenerative disease. Biophysics Reviews 5 (1), 011303 (2024), doi:10.1063/5.0180899, https://doi.org/10.1063/5.0180899.

16. T. B. Thompson, G. Meisl, T. P. J. Knowles, A. Goriely, The role of clearance mechanisms in the kinetics of pathological protein aggregation involved in neurodegenerative diseases. The Journal of Chemical Physics 154 (12), 125101 (2021), doi:10.1063/5.0031650, https://doi.org/10.1063/5.0031650.

17. T. J. Zwang, et al., Spatial characterization of tangle-bearing neurons and ghost tangles in the human inferior temporal gyrus with three-dimensional imaging. Brain Communications 5 (3), fcad130 (2023), doi:10.1093/braincomms/fcad130, https://doi.org/10.1093/braincomms/fcad130.

18. E. Dimou, et al., Super-resolution imaging unveils the self-replication of tau aggregates upon seeding. Cell Reports 42 (7), 112725 (2023), doi:10.1016/j.celrep.2023.112725, https://doi.org/10.1016/j.celrep.2023.112725.

19. T. Miller, J. Wu, S. Venkatesan, A. R. Gama, R. Halfmann, DAmFRET measures saturating concentrations and toxicities of protein phase transitions in vivo. Molecular Biology of the Cell 34 (6), br7 (2023), doi:10.1091/mbc.E22-11-0503, https://doi.org/10.1091/mbc.E22-11-0503.

20. C. Bellenguez, et al., New insights into the genetic etiology of Alzheimer’s disease and related dementias. Nature Genetics 54 (4), 412–436 (2022), doi:10.1038/s41588-022-01024-z, https://doi.org/10.1038/s41588-022-01024-z.

21. H. Koga, S. Kaushik, A. M. Cuervo, Protein homeostasis and aging: The importance of exquisite quality control. Ageing Research Reviews 10 (2), 205–215 (2011), doi:10.1016/j.arr.2010.02.001, https://doi.org/10.1016/j.arr.2010.02.001.

22. J. Keller, F. Huang, W. Markesbery, Decreased levels of proteasome activity and proteasome expression in aging spinal cord. Neuroscience 98 (1), 149–156 (2000), doi:10.1016/S0306-4522(00)00067-1, https://doi.org/10.1016/S0306-4522(00)00067-1.

23. T. P. J. Knowles, et al., An Analytical Solution to the Kinetics of Breakable Filament Assembly. Science 326 (5959), 1533–1537 (2009), doi:10.1126/science.1178250, https://doi.org/10.1126/science.1178250.

24. S. I. A. Cohen, M. Vendruscolo, C. M. Dobson, T. P. J. Knowles, Nucleated polymerization with secondary pathways. III. Equilibrium behavior and oligomer populations. The Journal of Chemical Physics 135 (6), 065107 (2011), doi:10.1063/1.3608918, https://doi.org/10.1063/1.3608918.

25. E. Fertan, et al., Super-resolution microscopy of alpha-synuclein aggregates in brain samples indicates a subset of cells have disrupted protein homeostasis (2025), doi:10.1101/2025.09.16.676632, https://doi.org/10.1101/2025.09.16.676632.

26. G. Meisl, et al., Molecular mechanisms of protein aggregation from global fitting of kinetic models. Nature Protocols 11 (2), 252–272 (2016), publisher: Nature Publishing Group, doi: 10.1038/nprot.2016.010, https://doi.org/10.1038/nprot.2016.010.

27. F. U. Hartl, A. Bracher, M. Hayer-Hartl, Molecular chaperones in protein folding and proteostasis. Nature 475 (7356), 324–332 (2011), doi:10.1038/nature10317, https://doi.org/10.1038/nature10317.

28. B. Chen, M. Retzlaff, T. Roos, J. Frydman, Cellular Strategies of Protein Quality Control. Cold Spring Harbor Perspectives in Biology 3 (8), a004374–a004374 (2011), doi:10.1101/cshperspect.a004374, http://doi.org/10.1101/cshperspect.a004374.

29. T. Khan, et al., Quantifying Nucleation In Vivo Reveals the Physical Basis of Prion-like Phase Behavior. Molecular Cell 71 (1), 155–168.e7 (2018), doi:10.1016/j.molcel.2018.06.016, https://doi.org/10.1016/j.molcel.2018.06.016.

30. T. M. Dawson, T. E. Golde, C. Lagier-Tourenne, Animal models of neurodegenerative diseases. Nature Neuroscience 21 (10), 1370–1379 (2018), doi:10.1038/s41593-018-0236-8, https://doi.org/10.1038/s41593-018-0236-8.

31. A. Abdelgawad, et al., Predicting longitudinal brain atrophy in Parkinson’s disease using a Susceptible-Infected-Removed agent-based model. Network Neuroscience 7 (3), 906–925 (2023), doi:10.1162/netna00296, https://doi.org/10.1162/netn_a_00296.

32. E. Head, W. Silverman, D. Patterson, I. T. Lott, Aging and Down Syndrome. Current Gerontology and Geriatrics Research 2012, 412536 (2012), doi:10.1155/2012/412536, https://doi.org/10.1155/2012/412536.

33. B. Rumble, et al., Amyloid A4 Protein and Its Precursor in Down’s Syndrome and Alzheimer’s Disease. New England Journal of Medicine 320 (22), 1446–1452 (1989), doi: 10.1056/NEJM198906013202203, http://doi.org/10.1056/NEJM198906013202203

34. W. B. Zigman, I. T. Lott, Alzheimer’s disease in Down syndrome: Neurobiology and risk. Mental Retardation and Developmental Disabilities Research Reviews 13 (3), 237–246 (2007), eprint: https://onlinelibrary.wiley.com/doi/pdf/10.1002/mrdd.20163, doi:10.1002/mrdd.20163, https://doi.org/10.1002/mrdd.20163.

35. A. B. Singleton, et al., α-Synuclein Locus Triplication Causes Parkinson’s Disease. Science 302 (5646), 841–841 (2003), publisher: American Association for the Advancement of Science, doi:10.1126/science.1090278, https://doi.org/10.1126/science.1090278.

36. M.-C. Chartier-Harlin, et al., α-synuclein locus duplication as a cause of familial Parkinson’s disease. The Lancet 364 (9440), 1167–1169 (2004), doi:10.1016/S0140-6736(04)17103-1, https://doi.org/10.1016/S0140-6736(04)17103-1.

37. L. V. C. Miller, et al., Tau assemblies do not behave like independently acting prion-like particles in mouse neural tissue. Acta Neuropathologica Communications 9 (1), 41 (2021), doi:10.1186/s40478-021-01141-6, https://doi.org/10.1186/s40478-021-01141-6.

38. B. J. Tuck, et al., Cholesterol determines the cytosolic entry and seeded aggregation of tau. Cell Reports 39 (5), 110776 (2022), doi:10.1016/j.celrep.2022.110776, https://doi.org/10.1016/j.celrep.2022.110776.

39. E. Fertan, et al., Clearance of beta-amyloid and tau aggregates is size dependent and altered by an inflammatory challenge. Brain Communications 7 (1), fcae454 (2024), doi:10.1093/braincomms/fcae454, https://doi.org/10.1093/braincomms/fcae454.

40. D. R. Whiten, et al., Tumour necrosis factor induces increased production of extracellular amyloid-β- and α-synuclein-containing aggregates by human Alzheimer’s disease neurons. Brain Communications 2 (2), fcaa146 (2020), doi:10.1093/braincomms/fcaa146, https://doi.org/10.1093/braincomms/fcaa146.

41. A. Ahern, T. B. Thompson, H. Oliveri, S. Lorthois, A. Goriely, Modelling cerebrovascular pathology and the spread of amyloid beta in Alzheimer’s disease. Proceedings of the Royal Society A: Mathematical, Physical and Engineering Sciences 481 (2311) (2025), publisher: The Royal Society, doi:10.1098/rspa.2024.0548, https://doi.org/10.1098/rspa.2024.0548.

42. C. H. Van Dyck, et al., Lecanemab in Early Alzheimer’s Disease. New England Journal of Medicine 388 (1), 9–21 (2023), doi:10.1056/NEJMoa2212948, https://doi.org/10.1056/NEJMoa2212948.

43. M. Vaz, V. Silva, C. Monteiro, S. Silvestre, Role of Aducanumab in the Treatment of Alzheimer’s Disease: Challenges and Opportunities. Clinical Interventions in Aging Volume 17, 797–810 (2022), doi:10.2147/CIA.S325026, https://doi.org/10.2147/CIA.S325026.

44. E. Fertan, et al., Lecanemab preferentially binds to smaller aggregates present at early Alzheimer’s disease. Alzheimer’s & Dementia 21 (4), e70086 (2025), eprint: https://alzjournals.onlinelibrary.wiley.com/doi/pdf/10.1002/alz.70086, doi:10.1002/alz.70086, https://doi.org/10.1002/alz.70086.

45. E. N. Neumann, et al., Brainwide silencing of prion protein by AAV-mediated delivery of an engineered compact epigenetic editor. Science 384 (6703), ado7082 (2024), doi:10.1126/science.ado7082, https://doi.org/10.1126/science.ado7082.

46. A. L. Edwards, et al., Exploratory Tau Biomarker Results From a Multiple Ascending-Dose Study of BIIB080 in Alzheimer Disease: A Randomized Clinical Trial. JAMA Neurology 80 (12), 1344 (2023), doi:10.1001/jamaneurol.2023.3861, https://doi.org/10.1001/jamaneurol.2023.3861.

47. D. D. Sahtoe, et al., Design of amyloidogenic peptide traps. Nature Chemical Biology 20 (8), 981–990 (2024), doi:10.1038/s41589-024-01578-5, https://doi.org/10.1038/s41589-024-01578-5.

48. G. Meisl, et al., Differences in nucleation behavior underlie the contrasting aggregation kinetics of the Aβ40 and Aβ42 peptides. Proceedings of the National Academy of Sciences 111, 9384–9389 (2014), tex.eprint: http://www.pnas.org/content/early/2014/06/16/1401564111.full.pdf+html,doi:10.1073/pnas.1401564111, http://doi.org/10.1073/pnas.1401564111.

49. G. Meisl, X. Yang, C. M. Dobson, S. Linse, T. P. J. Knowles, Modulation of electrostatic interactions to reveal a reaction network unifying the aggregation behaviour of the Aβ42 peptide and its variants. Chemical Science 8 (6), 4352–4362 (2017), publisher: The Royal Society of Chemistry, doi:10.1039/C7SC00215G, http://doi.org/10.1039/C7SC00215G.

50. R. Frankel, et al., Autocatalytic amplification of Alzheimer-associated Aβ42 peptide aggregation in human cerebrospinal fluid. Communications Biology 2 (1), 365 (2019), doi: 10.1038/s42003-019-0612-2, https://doi.org/10.1038/s42003-019-0612-2.

51. P. Flagmeier, et al., Mutations associated with familial Parkinson’s disease alter the initiation and amplification steps of α-synuclein aggregation. Proceedings of the National Academy of Sciences 113 (37), 10328–10333 (2016), doi:10.1073/pnas.1604645113, https://doi.org/10.1073/pnas.1604645113.

52. X. Yang, et al., On the role of sidechain size and charge in the aggregation of Aβ42 with familial mutations. Proceedings of the National Academy of Sciences 115 (26) (2018), doi: 10.1073/pnas.1803539115, https://doi.org/10.1073/pnas.1803539115.

53. M. Kotas, R. Medzhitov, Homeostasis, Inflammation, and Disease Susceptibility. Cell 160 (5), 816–827 (2015), doi:10.1016/j.cell.2015.02.010, https://doi.org/10.1016/j.cell.2015.02.010.

54. T. A. Cole, et al., α-Synuclein antisense oligonucleotides as a disease-modifying therapy for Parkinson’s disease. JCI Insight 6 (5), e135633 (2021), doi:10.1172/jci.insight.135633, https://doi.org/10.1172/jci.insight.135633.

55. J. W. Smit, et al., Phase 1/1b Studies of UCB0599, an Oral Inhibitor of α-Synuclein Misfolding, Including a Randomized Study in Parkinson’s Disease. Movement Disorders 37 (10), 2045– 2056 (2022), doi:10.1002/mds.29170, https://doi.org/10.1002/mds.29170.

56. J. Sevigny, et al., The antibody aducanumab reduces Aβ plaques in Alzheimer’s disease. Nature 537 (7618), 50–56 (2016), doi:10.1038/nature19323, https://doi.org/10.1038/nature19323.

57. S. I. A. Cohen, et al., Nucleated polymerization with secondary pathways. I. Time evolution of the principal moments. The Journal of chemical physics 135 (6), 065105 (2011), publisher: Department of Chemistry, University of Cambridge, Lensfield Road, Cambridge CB2 1EW, United KingdomNanoscience Centre, University of Cambridge, J J Thomson Avenue, Cambridge CB3 0FF, United KingdomCavendish Laboratory, University of Cambridge, J J Thomson Avenue, Cambridge CB3 0HE, United Kingdom. tex.medline-pst: ppublish tex.owner: tk tex.timestamp: 2011.08.22, doi:10.1063/1.3608916, http://dx.doi.org/10.1063/1.3608916.

58. S. I. A. Cohen, M. Vendruscolo, C. M. Dobson, T. P. J. Knowles, Nucleated polymerization with secondary pathways. II. Determination of self-consistent solutions to growth processes described by non-linear master equations. The Journal of chemical physics 135 (6), 065106 (2011), publisher: Department of Chemistry, University of Cambridge, Lensfield Road, Cambridge CB2 1EW, United Kingdom. tex.medline-pst: ppublish tex.owner: tk tex.timestamp: 2011.08.22, doi:10.1063/1.3608917, http://dx.doi.org/10.1063/1.3608917.

59. S. I. A. Cohen, M. Vendruscolo, C. M. Dobson, T. P. J. Knowles, Nucleated polymerization with secondary pathways. III. Equilibrium behavior and oligomer populations. The Journal of chemical physics 135 (6), 065107 (2011), publisher: Department of Chemistry, University of Cambridge, Lensfield Road, Cambridge CB2 1EW, United Kingdom. tex.medline-pst: ppublish tex.owner: tk tex.timestamp: 2011.08.22, doi:10.1063/1.3608918, http://dx.doi.org/10.1063/1.3608918.

60. A. W. P. Fitzpatrick, et al., Cryo-EM structures of tau filaments from Alzheimer’s disease. Nature 547 (7662), 185–190 (2017), doi:10.1038/nature23002, https://doi.org/10.1038/nature23002.

61. J.-C. Yen, F.-J. Chang, S. Chang, A new criterion for automatic multilevel thresholding. IEEE Transactions on Image Processing 4 (3), 370–378 (1995), doi:10.1109/83.366472, https://ieeexplore.ieee.org/document/366472.

62. S. v. d. Walt, et al., scikit-image: image processing in Python. PeerJ 2, e453 (2014), publisher: PeerJ Inc., doi:10.7717/peerj.453, https://peerj.com/articles/453.

63. S. Venkatesan, T. S. Kandola, A. Rodríguez-Gama, A. Box, R. Halfmann, Detecting and Characterizing Protein Self-Assembly In Vivo by Flow Cytometry. Journal of Visualized Experiments (149), 59577 (2019), doi:10.3791/59577, https://app.jove.com/t/59577.

64. B. B. Holmes, et al., Proteopathic tau seeding predicts tauopathy in vivo. Proceedings of the National Academy of Sciences 111 (41), E4376–E4385 (2014), publisher: Proceedings of the National Academy of Sciences, doi:10.1073/pnas.1411649111, https://www.pnas.org/doi/10.1073/pnas.1411649111.

65. D. F. Berenson, E. Zatulovskiy, S. Xie, J. M. Skotheim, Constitutive expression of a fluorescent protein reports the size of live human cells. Molecular Biology of the Cell 30 (24), 2985–2995 (2019), doi:10.1091/mbc.E19-03-0171, https://www.molbiolcell.org/doi/10.1091/mbc.E19-03-0171.

66. Z. Liu, et al., Systematic comparison of 2A peptides for cloning multi-genes in a polycistronic vector. Scientific Reports 7 (1), 2193 (2017), doi:10.1038/s41598-017-02460-2, https://www.nature.com/articles/s41598-017-02460-2 “http://www.nature.com/articles/s41598-017-02460-2”.

67. G. Meisl, et al., Scaling behaviour and rate-determining steps in filamentous self-assembly. Chemical Science 8 (10), 7087–7097 (2017), publisher: The Royal Society of Chemistry, doi:10.1039/C7SC01965C, http://dx.doi.org/10.1039/C7SC01965C.

