## Supplementary Information for "Neurodegeneration emerges at a cellular tipping point between aggregate accumulation and removal"

### Contents

**1 Mathematical Model of Aggregation and Removal** **S4**

1.1 Model including primary/secondary nucleation, elongation and chaperone mediated removal. . . . . . . . . . . . . . . . . . . . . . . . . . . . . . . S4

1.1.1 Parameters used in the text. . . . . . . . . . . . . . . . . . . . . . . . . S5

1.2 Minimal model and Simulation. . . . . . . . . . . . . . . . . . . . . . . . . . . . S5

**2 Aggregate Size Distributions in the Presence of Removal** **S8**

2.1 Model of Size Distributions gives Geometric Decay . . . . . . . . . . . . . . . . . . S8

2.2 Aggregate Size Distributions with Two Cell Types . . . . . . . . . . . . . . . . . . . S9

2.3 Fitting of Measured Size Distributions . . . . . . . . . . . . . . . . . . . . . . . S9

2.4 Monomer conversion factor . . . . . . . . . . . . . . . . . . . . . . . . . . . . S10

**3 High-resolution Imaging of Aggregates from Human Samples** **S11**

3.1 Single-molecule pull-down for determination of Aggregate Size Distributions in Brain Homogenate . . . . . . . . . . . . . . . . . . . . . . . . . . . . . . S11

3.2 Direct Imaging in Brain Slices . . . . . . . . . . . . . . . . . . . . . . . . . . . S13

3.2.1 Optical Setups . . . . . . . . . . . . . . . . . . . . . . . . . . . . . S13

3.2.2 FFPE Human Brain Slices . . . . . . . . . . . . . . . . . . . . . . . . S13

3.2.3 Puncta Detection from Raw Brain Images . . . . . . . . . . . . . . . . S14

**4 Quantifying critical monomer concentration with DAmFRET** **S14**

4.1 Distributed Amphifluoric FRET . . . . . . . . . . . . . . . . . . . . . . . . . . S14

4.1.1 Yeast experiments . . . . . . . . . . . . . . . . . . . . . . . . . . . S19

4.1.2 Tau amyloid biosensor cells . . . . . . . . . . . . . . . . . . . . . . . S19

4.1.3 Human cell culture and plasmid transfections . . . . . . . . . . . . . . S20

4.1.4 DAmFRET data collection for HEK293T cells . . . . . . . . . . . . . . S20

4.2 Determining thresholds to describe cellular states. . . . . . . . . . . . . . . . . . S21

4.2.1 Distinct aggregate populations (main text Fig. 3) . . . . . . . . . . . . . S22

4.2.2 Response to preformed fibrils (main text Fig. 4) . . . . . . . . . . . . . S22

4.2.3 Response to designed aggregation inhibitors (main text Fig. 4) . . . . . . . S23

4.3 Fitting kinetic rates in the presence of inhibitors. . . . . . . . . . . . . . . . . . . . . . S25
4.4 Relating critical concentration to effective rates. . . . . . . . . . . . . . . . . . . . . . S26

**5 Effect of the Introduction of Preformed Seeds** **S27**
5.1 Multi-hits in a Poisson Process . . . . . . . . . . . . . . . . . . . . . . . . . . . . . S27
5.2 Fitting of Seeding Data . . . . . . . . . . . . . . . . . . . . . . . . . . . . . . . . . S28

### 1 Mathematical Model of Aggregation and Removal

#### 1.1 Model including primary/secondary nucleation, elongation and chaperone mediated removal.

We derive the equation that describes the evolution of the fraction of aggregated protein, $\mu$. Here, we briefly overview a specific example of the aggregation and removal kinetics, but discuss this in further detail and explore generalisations in *Auxiliary Supplementary Materials Cotton et al. 2025*. We consider an individual, well mixed cell, containing a population of aggregates with a concentration of $f(i)$ aggregates of length $i$ and a concentration of free monomeric protein, $m$. We use established mechanisms to determine how this population of aggregates evolves in time. We consider a specific realisation of an aggregating system where aggregates can be nucleated via a spontaneous nucleation process to produce aggregates of size $n_c$ or via a catalysed secondary nucleation process that produces aggregates of size $n_2$. The secondary nucleation is catalysed by existing aggregates. We additionally allow aggregates to elongate via a polymerisation process: aggregates can grow from size $i \longrightarrow i+1$ via additional monomers attaching to either end. To extend this model to cell systems, we also consider removal mechanisms of an aggregate. The master equation that describes the evolution of an aggregate concentration of $f(i)$ in this systems is

$$\frac{\mathrm{d}f(i)}{\mathrm{d}t} = \delta_{i,n_C} k_n m^{n_C} + 2k_{\mathrm{on}} m (f(i-1) - f(i)) + \delta_{i,n_C} k_2 m^{n_2} \left( \sum_{i=2}^{\infty} f(i) \right) - r_i \tag{S1}$$

where the rate constants $k_n$, $k_2$ and $k_+$ correspond to the primary nucleation, secondary nucleation and polymerisation processes respectively. The removal rate of aggregates of size $i$ is $r_i$.

These dynamics produce a closed system of moment equations that define the distribution of aggregates. We define the total mass of aggregates $M = \sum_i i f(i)$ and the total number of aggregates, $P = \sum_i f(i)$. Summing over equation (S1) gives the evolution of the aggregate mass as

$$\frac{\mathrm{d}M}{\mathrm{d}t} = 2k_+ mP + n_c k_n m^{n_c} + n_2 k_2 m^{n_2} M - \sum_i^{\infty} i r_i. \tag{S2}$$

We further reduce the system by connecting $M$ and $P$ using the average aggregate length, $\bar{l}$. Prior work (*57–59*) has shown that as a system transitions to a runaway aggregation state, $\bar{l} = \sqrt{2k_+/k_2 m_0}$ and so we use this to connect the aggregate number and mass concentrations: $P = M/\bar{l}$. Additionally we assume that the removal mechanisms follow chaperone mediated kinetics with a chaperone that is shared between aggregates and that binds to and breaks down aggregates of all sizes with the same rates. The total removal of aggregate mass is therefore

$$\sum_i i r_i = \frac{\lambda M}{1 + P/K_\lambda} = \frac{\lambda M}{1 + M/\bar{l}K_\lambda}. \tag{S3}$$

Combining all of these ingredients we find a reduced model of the evolution of the aggregate mass as

$$\frac{\mathrm{d}M}{\mathrm{d}t} = 2k_+ m M/\bar{l} + n_c k_n m^{n_c} + n_2 k_2 m^{n_2} M - \frac{\lambda M}{1 + M/\bar{l}K_\lambda}. \tag{S4}$$

We assume that cells regulate the total protein concentration and so we set $m_{\text{tot}} = m_0 + M$. We define the fraction of aggregated cells as $\mu = M/m_{\text{tot}}$ and thus we get

$$\begin{aligned}\frac{\mathrm{d}\mu}{\mathrm{d}t} &= n_{\text{c}} k_{\text{n}} (1-\mu)^{n_{\text{c}}} m_{\text{tot}}^{n_{\text{c}}-1} + n_2 k_2 \mu (1-\mu)^{n_2} m_{\text{tot}}^{n_2} \\ &\quad + 2k_{\text{on}} \mu (1-\mu) m_{\text{tot}}/\bar{l} - \frac{\lambda \mu}{1 + \mu m_{\text{tot}}/(K_\lambda \bar{l})},\end{aligned} \tag{S5}$$

##### 1.1.1 Parameters used in the text.

Below we outline the parameter values used to draw the stable and unstable branches in each figure (using equation (S5)):

- **Fig. 3:** (c) values in Table S1. For (d) see Section 1.2.
- **Fig. 4:** (b) values in Table S1 (c) *Normal* stability line uses values in Table S1, *Stressed* uses values as in Table S1, but with $\lambda = 0.05 \times 10^4 \text{hr}^{-1}$.
- **Fig. 5:** (b) For the *No Inhibition* plot, values in Table S1. For the *Inhibition* plot, values in Table S1, except $k_2 = 1.2 \times 10^{12} \text{M}^{-n_2} \text{hr}^{-1}$.

#### 1.2 Minimal model and Simulation.

To simulate the DAmFRET experimental data, we create a minimal model of aggregation, inspired by equation (S4). The majority of the phase plot is dominated by the balance between removal and self-replication, with the primary nucleation process being important only for the position of the stable lower branch. Since in most experimental data, including the DAmFRET experiments, the low aggregate concentrations in the lower branch cannot be resolved from the noise, we can simplify the description by omitting this nucleation process, effectively setting the lower stable branch to $M = 0$, without a significant sacrifice in the accuracy of the description. We assume $n_2 = 1$, so that we can write equation (S4) as

$$\frac{\mathrm{d}M}{\mathrm{d}t} = kM(m_{\mathrm{tot}} - M) - \frac{\lambda M}{1 + M/\tilde{K}}. \tag{S6}$$

where $k = (2k_+/\bar{l} + n_2 k_2)$, $\tilde{K} = \bar{l} K_\lambda$ and we use the total protein mass concentration $m_{\mathrm{tot}}$ as this is experimentally accessible.

The $M = 0$ steady state becomes unstable when $m_{\mathrm{tot}} > \lambda/k$, which is the critical monomer concentration. We use this value from experiments at 40hr incubation, 0.85 a.u., and set $\lambda/k = 0.85$. Since equation (S6) is quadratic in $M$, we can solve exactly for the steady states to get

$$M^*_\pm = \frac{1}{2}\left(-\tilde{K} + m_{\mathrm{tot}} \pm \sqrt{(\tilde{K} + m_{\mathrm{tot}})^2 - 4\tilde{K}\lambda/k}\right) \tag{S7}$$

where the positive root is stable and the negative root is unstable. We additionally need to choose $\tilde{K}$ and we arbitrarily set this to be $\tilde{K} = \lambda/3k = 0.37$. Evaluating $M^*$ gives the fixed point lines shown in Fig. 1(d) in the main text (the $y$-axis in the plot is $M^*/m_{\mathrm{tot}}$).

Additionally we can simulate data where the parameters in equation (S6) are drawn from some distribution. We generate this data as follows. Firstly we take every recorded value of $m_{\mathrm{tot}}$ simulated and compare it to a randomly generated value of $\lambda/k$, drawn from a normal distribution with mean 0.85 a.u and a standard deviation of 0.25 (used as this is half the distance between the protein concentration at which 25% and 75% of the cells are in an aggregated state). If the measured $m_{\mathrm{tot}}$ is below the randomly generated value, then we set the steady state to $M^* = 0$. If not, we draw a random value of $\tilde{K}$ from a normal distribution with mean 0.37 and an arbitrarily chosen width of 0.05, and use these values to evaluate equation (S7) to determine $M^*$. To simulate the real data collection process we add a randomly generated noise, to give a measured FRET signal, $M_m$. We model this as having a proportional and constant offset measurement noise, such that $M_m = \eta_1 M^* + \eta_2$, where $\eta_1$ is drawn from a normal distribution of mean 1, standard deviation 0.2 and $\eta_2$ is drawn from a normal distribution of mean 0, standard deviation 0.01. The plot shows $M^*/m_{\mathrm{tot}}$ against $m_{\mathrm{tot}}$.

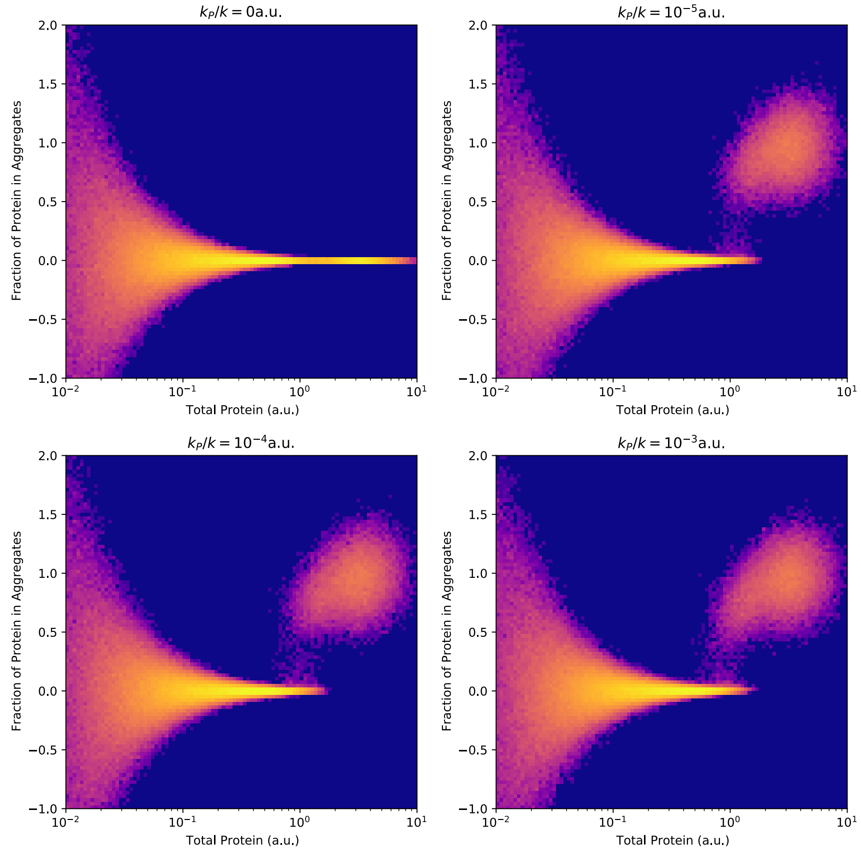


**Figure S1**: Simulated data of a model with nucleation as described in equations (S8).

We can additionally include a primary nucleation term in the minimal model given by equation (S6) to give

$$\frac{\mathrm{d}M}{\mathrm{d}t} = kM(m_{\mathrm{tot}} - M) - \frac{\lambda M}{1 + M/\tilde{K}} + k_P(m_{\mathrm{tot}} - M). \tag{S8}$$

This updated model can also be used to simulate the aggregation phase plane given by Fig. 2a in the main text. For every DAmFRET measurement, we initialise a system with $m_{tot}$ as measured by acceptor fluorescence and with $M = 0.01$. We evolve each cell using equation (S8) where $k$ and $\tilde{K}$ are drawn from the same distribution as before and we choose a the same value of $k_p/k$ for all cells ($k_p/k = 0$a.u., $10^{-5}$a.u., $10^{-4}$a.u., $10^{-3}$a.u.). This range of values is based on the condition that the system is dominated by self-replication (*14*), thus requiring that $k_p/k \ll m_{tot}$. We evolve each system until the steady state distribution is reached and then plot the 2d histogram as from the DAmFRET measurement ($M/m_{tot}$ vs $m_{tot}$).

### 2 Aggregate Size Distributions in the Presence of Removal

#### 2.1 Model of Size Distributions gives Geometric Decay

The majority of changes in aggregate size will be from aggregates growing by the addition of a monomeric protein. This elongation process happens at some rate $k_+ m_0$ where $k_+$ is the rate constant for polymerisation and $m_0$ is the monomer protein concentration. When the system is in a quasi-steady size distribution this elongation process will be balanced by removal via some per aggregate removal rate, $r$. Note this is an instantaneous removal rate for an individual aggregate and we ignore any functional dependence on this rate, in comparison to Section 1.1, this would be $r = \lambda/(1 + M/\bar{l}K_\lambda)$. When the aggregates are larger than typical nucleation sizes and the depolymerisation rate is small, these two processes will dominate the size distribution of aggregates. This model is applicable only within a specific range of aggregate sizes: aggregates must be (1) sufficiently large to be unaffected by nucleation-related fluctuations or conformational changes, but (2) not so large that alternative removal mechanisms are involved.

The system dynamics will establish a steady state aggregate distribution. The system shown in Fig. S2 has a constant distribution when $k_+ m_0 f(i-1) = (k_+ m_0 + r) f(i)$. We can therefore write the concentration of aggregates of some size $i$ to be $f(i) \propto \alpha^i$, with

$$\alpha = \frac{1}{\frac{r}{k_+ m_0} + 1}. \tag{S9}$$

Changes to overall nucleation rate will simply modify the constant of proportionality in the probability distribution but not affect $\alpha$. On a log-linear histogram showing the frequency of aggregates at every size, the geometric distribution will appear as a straight line. The slope of this line will be given by $log_{10}(\alpha)$ (when working in base 10 logarithms).

The ratio of the removal to elongation rate, $r/(k_+ m_0)$, can be calculated given $\alpha$ by inverting equation (S9) to give $r/(k_+ m_0) = \alpha^{-1} - 1$. We can define the *normalised removal*, $\tilde{r} = r/(k_+ m_0)$ and compare this across different aggregate populations to compare the balance of elongation and removal mechanisms generating the aggregate populations. This model holds as long as the number of growth sites is constant and independent of aggregate size.

The aggregate size distribution is fit between a minimum and maximum aggregate size, $i_{min}$ and $i_{max}$ respectively. This is motivated by the discussion above, however often we set $i_{max}$ to

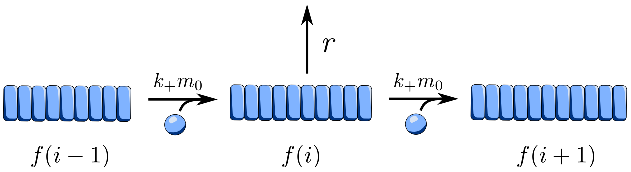


**Figure S2**: Geometric growth schematic.

prevent individual outliers at very large aggregate sizes from biasing the fits. We therefore have a normalised aggregate size distribution, where the probability of an aggregate being size $i$ is

$$p_i = \frac{1-\alpha}{\alpha^{i_{min}} - \alpha^{i_{max}+1}}\alpha^i. \tag{S10}$$

#### 2.2 Aggregate Size Distributions with Two Cell Types

When some of the cells are in the pathological state, we expect that the size distribution will be made up of aggregates in both healthy and in pathological cells. The distribution therefore looks like

$$p_i = (1-\beta)p_i^h + \beta p_i^d \tag{S11}$$

where $p_i^h$ ($p_i^d$) is the normalised aggregate size distribution in healthy (pathological) cells, which is simply given by equation (S10) using the decay factor, $\alpha_H$ ($\alpha_D$). The variable $\beta$ is the fraction of cells in the tissue that are in the pathological state.

The two different decay factors mean that we can compare the normalised removal in health and disease and define the ratio of the normalised removal between health and disease, $r_C$, as

$$r_C = \frac{\tilde{r}_d}{\tilde{r}_h} = \frac{\alpha_D^{-1} - 1}{\alpha_H^{-1} - 1}. \tag{S12}$$

#### 2.3 Fitting of Measured Size Distributions

We use a two-step process to fit the aggregate size distributions. Firstly, we fit equation S10 for the healthy control samples, to determine the geometric decay factor in health, $\alpha_H$. The results of this inference are shown in Table S2. This is the fit shown for the control aggregates sizes.

Secondly, we fix this value of $\alpha_H$ using the maximum likelihood values from the control fits and we infer $\beta$ and $r_c$. We do this by constructing a likelihood of the observed aggregate size distribution as the product of the probabilities of each aggregate, using equation (S11) (alongside equations (S9) and (S12)). We use a uniform prior for $\beta$ between 0 and 1, and for $r_C$ we assume a uniform prior between 0 and 1. The results of this inference are shown in Table S3.

#### 2.4 Monomer conversion factor

For aggregate length data by super-resolution microscopy, we convert the aggregate sizes to monomers by multiplying the length of the aggregate in nanometres by 4, from the assumption of a double stranded aggregate with a beta-sheet separation of 0.5 nm (*60*). However, as we now demonstrate, the relative removal rates extracted are robust with respect to the precise conversion factor used to relate physical size to monomer number.

Suppose we measure the relative removal between two systems, $A$ and $B$, resulting in the following ratio:

$$r_C = \frac{\tilde{r}_A}{\tilde{r}_B} = \frac{\alpha_A^{-1} - 1}{\alpha_B^{-1} - 1}. \tag{S13}$$

We define $\alpha = 1 - \epsilon\delta$, with $\delta \sim O(1)$. Substituting this into equation (S13), we find

$$\frac{\tilde{r}_A}{\tilde{r}_B} = \frac{\alpha_A^{-1} - 1}{\alpha_B^{-1} - 1} = \frac{\delta_A}{\delta_B}. \tag{S14}$$

Now consider the effect of an incorrect conversion from physical size to monomer number, represented by a constant scaling factor $q$. Under this error, the decay parameter is misestimated as $\tilde{\alpha} = \alpha^q$ leading to an incorrect estimate of the normalised removal ratio:

$$\frac{\tilde{r}_A}{\tilde{r}_B} = \frac{\tilde{\alpha}_A^{-1} - 1}{\tilde{\alpha}_B^{-1} - 1} = \frac{\alpha_A^{-q} - 1}{\alpha_B^{-q} - 1}. \tag{S15}$$

However, as aggregates are typically $\sim$ 100s of monomers long, we expect $\alpha \approx 1$ and $\epsilon \ll 1$. Expanding the misestimated normalised removal ratio in $\epsilon$, we find

$$\frac{\alpha_A^{-q} - 1}{\alpha_B^{-q} - 1} = \frac{(1 - \epsilon\delta_A)^{-q} - 1}{(1 - \epsilon\delta_B)^{-q} - 1} = \frac{\delta_A}{\delta_B} + \frac{(q-1)\delta_A(\delta_A - \delta_B)}{2\delta_B}\epsilon + O(\epsilon^2). \tag{S16}$$

To leading order in $\epsilon$, this expression reproduces the exact ratio in Eq. (S14). Therefore, the measurement of the ratio of the normalised removal is robust to errors in the absolute conversion from physical size to monomer count.

### 3 High-resolution Imaging of Aggregates from Human Samples

#### 3.1 Single-molecule pull-down for determination of Aggregate Size Distributions in Brain Homogenate

Post-mortem brain tissue from donors with AD, PSP, PiD (Picks Disease), CBD, PD, and neurologically healthy controls was obtained from the Cambridge and Edinburgh Brain Banks under approved ethical protocols (REC 16/WA/0240 and REC 21/ES/0087). Written informed consent from the donors and/or their next of kin was provided as appropriate. Donor characteristics are summarised in Tables S4 and S5. Tissue samples were processed as previously described (*10*).

In brief, the PD samples were soaked by cutting them into 300-350 mg pieces and placing them in 1.5 mL low-binding Eppendorf tubes (Thermo Scientific, Cat. 90410). 600 µl artificial CSF buffer (bio-techne TORIS, Cat. 3525) containing 150 nM Na+, 3 nM K+, 1.4 nM Ca2 +, 0.8 nM Mg2 +, 1 nM P and 155 nM Cl- with Halt™ protease and phosphatase inhibitor single-use cocktail (100x; Thermo Fischer Scientific, Cat. 78442) was added to each sample and incubated for 45-minutes at 4°C on a HulaMixer ™ (Thermo Scientific, Cat. 15920D). Then the samples were centrifuged at 17,000 G for 120-minutes at 4°C, the supernatant was collected and stored in a -80°C freezer until analysis. Meanwhile, the AD, PSP, PiD, CBD cases were homogenised by placing 120 mg of tissue in 1.5 mL low-binding Eppendorf tubes and adding 1 mm zirconium beads (Scientific Labs, Cat. SLS1414) and initially adding 1.2 ml artificial CSF with protease/phosphatase inhibitors and processing the samples on an electronic tissue homogeniser (VelociRuptor V2 Microtube Homogeniser, Scientific Labs, Cat. SLS1401), at 5 meters/sec for 2 cycles of 15 seconds, with a 10 second gap in between, followed by centrifugation at 17,000 G for 120-minutes at 4°C. The supernatant was collected and stored on wet ice, while the pallet was once again homogenised by adding 600 µl artificial CSF. The supernatants were mixed and stored in a -80°C freezer until analysis.

Single-molecule pull-down (SiMPull) and DNA-PAINT experiments were performed as described in Böken et al. (*11*) and Fertan et al. (*25*), using PEG functionalised glass coverslips and biotinylated monoclonal anti-tau (AT8 Invitrogen, MN1020b, HT7 Invitrogen MN1000b) and anti $\alpha$-synuclein (4B12, Thermo Scientific, MA1-90346) to capture the tau and $\alpha$-synuclein aggregates.

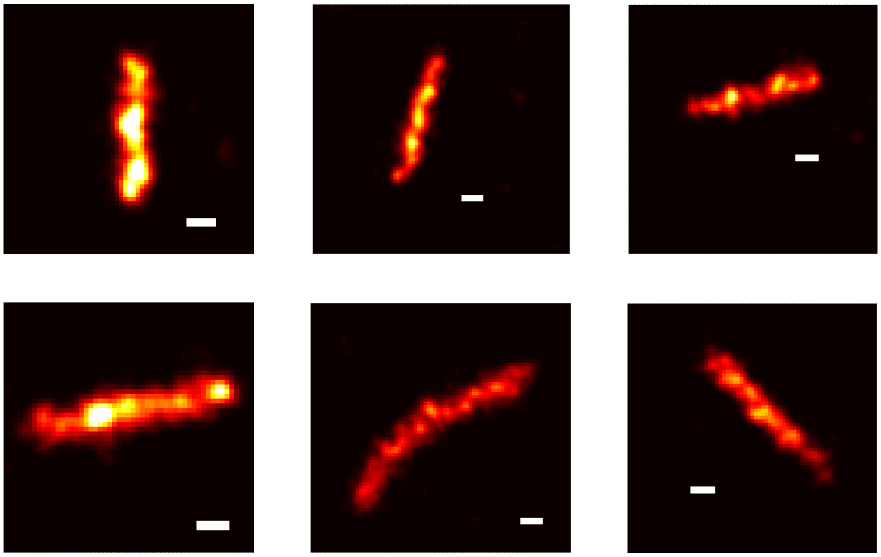

**Figure S3**: Exemplar images of fibrillar aggregates from single molecule super-resolution images. Scale bar, 100 nm.

For fluorescence imaging, samples were labelled with the same respective monoclonal DNA-conjugated antibodies (AT8 Invitrogen, MN1020, HT7 Invitrogen MN1000, 4B12, Thermo Scientific, MA1-90346, conjugated to DBCO TEG-AAACCACCACCACCACCACCACCACCACCACCA, ATDBio), and the corresponding atto655-conjugated imaging strand (TGGTGGT-AminoC7-atto655) was added. Imaging was conducted on a custom-built TIRF microscope using 638nm laser excitation for tau and 568 nm excitation for $\alpha$-synuclein. For each well, 4 field of views were collected in a 2-by-2 grid using an automated script (Micro-Manager) to avoid any bias in the selection. Images were acquired for 8000 frames of 100 ms exposure. The tau data used in this work was generated with the HT7 antibody.

Super-resolution reconstructions were performed as described in Böken et al. (*11*) using Picasso. Localisations were drift-corrected, filtered for precision (<30 nm), and analysed using DBSCAN clustering (radius of 0.3 ($\approx$35nm) and minimum density of 5) and skeletonised to quantify aggregate length. Fig. S3 shows examples of the aggregates' fibrillar shape and justifies the measurement of aggregate length.

#### 3.2 Direct Imaging in Brain Slices

##### 3.2.1 Optical Setups

Data shown in main text Fig. 1 was generated by direct imaging of tissue samples in situ.

Experiments were performed on one of two microscopes: a widefield single-molecule microscope (herein called 'Microscope 1') or a spinning-disk confocal microscope ('Microscope 2'). These have been described before, as 'Microscope 1' and 'Microscope 3' in Fu et. al. (*13*), respectively.

Microscope 1 is a widefield fluorescence microscope equipped with a 488 nm and a 561 nm laser, with excitation of the samples performed at HILO illumination. The filtered fluorescence light was expanded (1.5x) and projected onto an electron-multiplying charge-coupled device (EMCCD, Evolve 512 Delta, Photometrics) operating in frame transfer mode with an electron multiplication gain of 250 ADU/photon.

Microscope 2 is a commercial spinning-disk confocal microscope (3i intelligent imaging) equipped with 488 (LuxX) and 561 (OBIS) nm lasers. The fluorescence was detected using one of two sCMOS cameras (Prime 95B, Teledyne Photometrics).

##### 3.2.2 FFPE Human Brain Slices

FFPE human brain slices were stained in accordance with the protocol laid out in Andrews et. al. (*12*) and in Fu et. al (*13*). Formalin-fixed paraffin-embedded (FFPE) tissue sections were obtained from the cingulate cortex and cut to 8 micron thickness. FFPE sections were baked at 37 degrees C for 24 hours followed by 60 degrees C overnight. Sections were deparaffinized in xylene, and rehydrated using graded alcohols. Non-specific binding was blocked with 1% bovine serum albumin (BSA) solution in PBS for 30 minutes. The tissue was then pressure cooked in citrate buffer at pH 6 for 10 minutes. Tissue sections were incubated with primary antibodies; anti-phosphorated $\alpha$-synuclein (ab184674, Abcam, 1:500; ab59264, Abcam, 1:200) ; Microtubule-Associated Protein 2 (ab254143, Abcam, 1:500); ionized calcium-binding adapter molecule 1 (Wako – 019-19741, FujiFilm, 1:1000) for 1h at room temperature. The sections were then washed three times for five minutes in PBS followed by the corresponding AlexaFluor secondary antibodies (anti-mouse 568—A11031, Thermo Fisher, anti-rabbit 568—A11011, Thermo Fisher, anti-mouse 488—A11001,

Thermo Fisher, anti-rabbit 488—A11008, Thermo Fisher, all at 1:200) for an additional hour at room temperature in the dark. Sections were then washed three times for five minutes again in PBS and incubated in Sudan Black (0.1% for 10 minutes, 199664-25G, Sigma Aldrich). Removal of Sudan Black occurred with three washes in 30% ethanol (E7148-500ML, Sigma Aldrich) before mounting with Vectashield+ (Vector Labs, H-1900) and coverslipping (VWR, 50 mm x 24 mm #1 thickness, Catalogue Number 48404-453) for imaging. Sections were stored at 4 degrees C until imaging was completed.

##### 3.2.3 Puncta Detection from Raw Brain Images

Aggregate detection proceeded using the RASP pipeline described in Fu et al. (*13*). To briefly summarise, images underwent a high-pass kernel, obtained through the difference between the original image and a Gaussian-blurred image ($\sigma$ = 1.4px), followed by a Laplacian-of-Gaussian (LoG) kernel ($\sigma$ = 2px). Puncta were selected as pixels in the top 95th percentile of brightness, and these were then accepted or rejected based on their integrated gradients and flatness (*13*). See Fig. S4 for exemplar puncta.

For cell mask detection, each 2D image was enhanced using a difference-of-Gaussian filter, with $\sigma_1$ = 2px and $\sigma_2$ = 60px. The image was then thresholded into a binary mask using a Yen threshold (*61*). This binary mask then had a binary opening operation applied to it with a disk morphology of radius 1 pixel, which was then followed by a binary closing operation with a disk morphology of radius 5 pixels. When all of the images in a volumetric stack had binary masks generated in this way, the scikit.image (*62*) function binary_fill_holes was applied to the full 3D volume, after which small holes of $<$ 100 voxels were removed. Post this, 3D objects below a specified cell size were removed to end up with the final cell masks used for analysis.

### 4 Quantifying critical monomer concentration with DAmFRET

#### 4.1 Distributed Amphifluoric FRET

In this work we take advantage of the high throughput Distributed Amphifluoric FRET assay (*29*). We use this assay to explore the phase plane of aggregation by sampling a large range of cellular

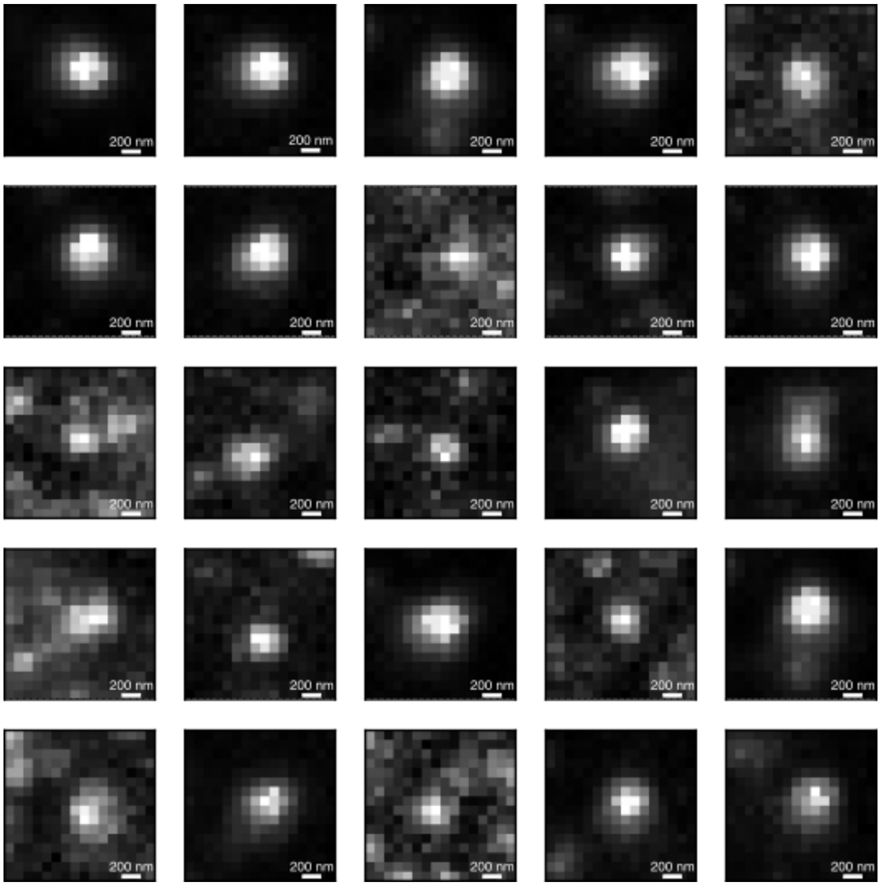


**Figure S4**: Gallery of example puncta detected as $\alpha$-synuclein oligomers using pS129 stain.

protein expression levels and measuring the fraction of aggregated protein after some incubation time. The enumerated list below provides details of the specific assay used in each figure. Raw DAmFRET data that support the findings of this study are openly available in Stowers Open Data Repository at `https://www.stowers.org/research/publications/LIBPB-2570`. We normalise the colours of the 2d histograms logarithmically and set 0 to be the dark background colour (equivalent colour to value 1). The following subsections explain the specific experimental setups.

1. **Main text Fig3** Expression of HET-s 218-289 in rhy2216 yeast cells. The x-axis parameter is acceptor fluorescence divided by the side scatter (SSC), a proxy for cell volume, thereby reporting the protein concentration in each cell (*19*). This allows us to compare concentrations across time points. On the y-axis we report "AmFRET", which is defined as FRET signal normalised by direct-excited acceptor fluorescence, to give a fraction of aggregated protein in the cell. This is correct up to a constant of proportionality from converting the signal to concentrations. N=3 for each time point and Fig. S5 shows consistent behaviour across all replicates.

2. **Main text Fig. 4** We treated our HEK293T cell-based tau biosensor line with various amounts of sonicated commercial pre-formed fibrils and incubated them for 48hrs. The relevant quantities are the acceptor fluorescence (x-axis), and the ratio of amphifluoric FRET over acceptor fluorescence (y-axis). We determined the fraction of aggregated cells under identical incubation and plot how these vary across dosages. N=2 and both replicates are shown in Fig. S8 for repeats.

3. **Main text Fig5** HEK293T cells expressing A$\beta$42 can also be engineered to express a designed peptide that inhibits aggregation (*47*) by means of a tandem ribosome skipping motif in the same open reading frame. This was the experimental setup used in Fig. 5 in the main text. The x-axis is the acceptor fluorescence and the y-axis is the ratio of amphifluoric FRET over acceptor fluorescence. N=3 and Fig. S6 shows consistent behaviour across all replicates.

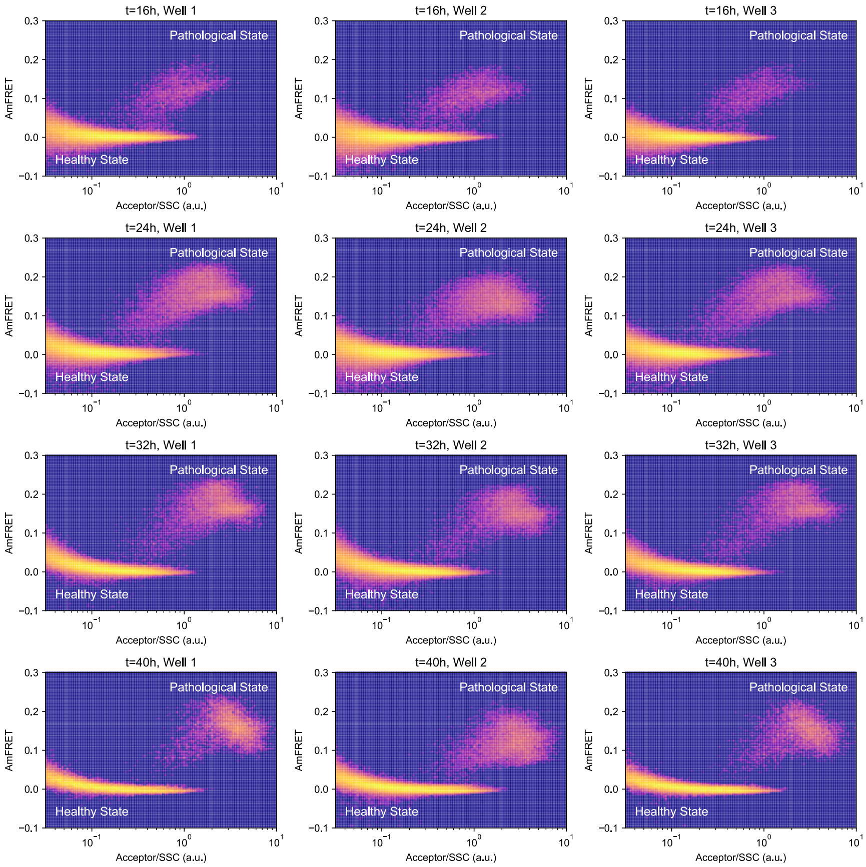


**Figure S5**: Experimental repeats for the DAmFRET data in main text Fig. 3. Three distinct populations were measured at every time.

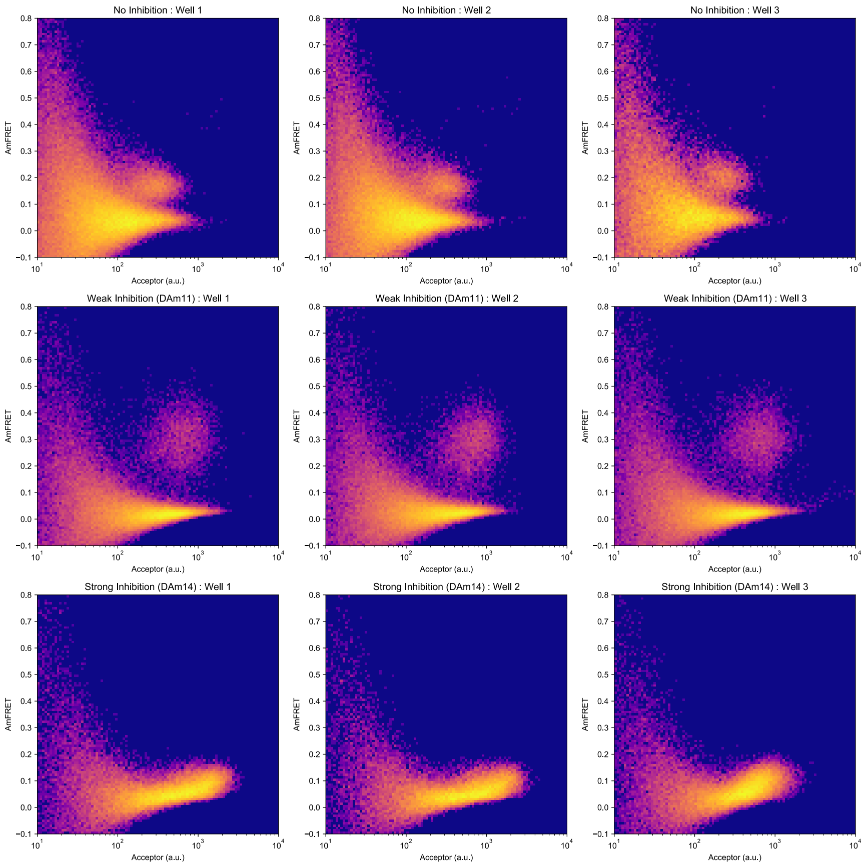


**Figure S6**: Experimental repeats for the DAmFRET data in main text Fig. 4. Three distinct populations were measured for each binder and the control.

##### 4.1.1 Yeast experiments

The yeast strain used (rhy2216) was generated by mating rhy1713 (*29*) with the NAM7 deletion strain from the MATa systematic deletion collection (Open Biosystems) and sporulating the resulting diploid to obtain a sporulant with the genotype MATa cln3Δ0::GAL1pr_WHI5_hphMX nam7Δ0::kanMX can1Δ::STE2pr_SpHIS5 lyp1Δ his3Δ1 leu2Δ0 met17Δ0 ura3Δ0. The NAM7 deletion was introduced to minimize degradation of ectopically expressed mRNA. Details of the plasmid (rhx0952), yeast manipulations, and yeast DAmFRET data collection, are as previously described (*19, 63*).

##### 4.1.2 Tau amyloid biosensor cells

A clonal tau amyloid biosensor cell line inspired by Holmes et al. (*64*), but using AmFRET instead of dual fluorophore FRET, was generated by integrating via lentivirus, plasmid rhm0523 into HEK293T. This plasmid was made by replacing the full open reading frame of CSII-prEF1a-mCherry-3xNLS (a gift from Jan Skotheim; Addgene plasmid # 125262; (*65*)) with an open reading frame encoding tau 2N4R 246-378 P301S fused at its C-terminus to a rigid linker, 4x(EAAAR), and mEos3.2, placing it under the control of the EF1$\alpha$ promoter. A clone was isolated with uniform mEos3.2 expression.

The HEK293T tau biosensor cells were cultured in DMEM medium warmed to 37°C supplemented with 10% FBS and 1% Pen-Strep. For a 12-well plate, 1mL of media was added per well. Cells were maintained in a humidified incubator at 37°C with 5% $CO_2$. Two days prior to fibril transfection, the cells were plated at $2 \times 10^5$ cells/well. Pre-formed fibrils of tau 2N4R P301S (StressMarq SPR-329C) were diluted in Opti-MEM™ to 1mg/mL, then sonicated for 2 minutes (2 cycles of 30 seconds of sonication and 30 seconds of pause) in a Diagenode Bioruptor Pico (20-60kHz). The fibers were diluted in Opti-MEM™ to a final volume of 100$\mu$L per replicate. For transfections, Lipofectamine™ 3000 Transfection Reagent (7uL/replicate) and Opti-MEM™ (93uL/replicate) were mixed and added to the different fibril dilutions after a 5-minute incubation period. The Lipofectamine™ and fiber mixture were incubated for 20 minutes and then 200uL of the mixture was added per replicate well. After incubating for 10, 24, or 48 hours, cells were collected and fixed for photoconversion and data collection.

###### 4.1.3 Human cell culture and plasmid transfections

HEK293T cells were cultured in DMEM medium warmed to 37°C supplemented with 10% FBS and 1% Pen-Strep. For a 12-well plate, 1mL of media was added per well. Cells were maintained in a humidified incubator at 37°C with 5% $CO_2$. Cells were plated at $4 \times 10^5$ cells/well in a 12-well plate one day prior to plasmid transfection. For each transfection replicate, 2$\mu$g DNA was added to 22$\mu$L Opti-MEM™ and 3$\mu$L FUGENE HD, for a final volume of 25$\mu$L. The mixture was incubated for 15 minutes then added to the cells. Four hours after transfection, doxycycline was added at 5$\mu$g/mL. After 24 hours, cells were refreshed with new media and doxycycline (5ug/mL). After an additional 48 hours, cells were collected with TrypLE into a 96-well plate, then fixed with 4% paraformaldehyde at 37°C for 5 minutes, and finally, resuspended in phosphate-buffered saline (PBS) supplemented with 10mM of ethylenediamine tetraacetic acid (EDTA).

For A$\beta$ (1-42) DAmFRET, plasmid rhm0864 was created by replacing the full open reading frame in pCW57.1 (a gift from David Root; Addgene plasmid # 41393) with A$\beta$ (1-42) fused at its N-terminus to mEos3.2 and a rigid linker 4x(EAAAR), placing it under the control of the doxycycline-inducible "tight Tet-responsive element (TRE)" promoter. For plasmids co-expressing DAm11 or DAm14 with A$\beta$ (rhm0886 and rhm0888, respectively), the inhibitor-encoding sequences followed by two copies of the co-translational ribosome skipping sequence (P2A; (*66*)) were inserted at the N-terminus of mEos3.2 in rhm0864, allowing for stoichiometrically fixed expression of inhibitor with mEos3.2-fused A$\beta$, each as separate polypeptides. All plasmids and sequences are available upon request.

###### 4.1.4 DAmFRET data collection for HEK293T cells

Prior to cytometry, microplates containing fixed cells were illuminated with an OmniCure S2000 Elite UV lamp fitted with a 320–500 nm (violet) filter and a beam collimator (Exfo), positioned 45 cm above the plate, for a duration of 5 min to deliver an empirically determined total energy of 12.048 J/cm$^2$, while shaking at 1000 rpm with a 2 mm orbit diameter.

DAmFRET data were collected using a BioRad ZE5 cytometer with the detection channels 460/22 nm (405 nm laser), 488/10 nm (405 nm laser), 525/35nm (488 nm laser), 593/52nm (488 nm laser), and 589/15nm (561 nm laser) to measure the signals for Autofluorescence, Side Scatter

(SSC) and Forward Scatter (FSC), Donor, FRET, and Acceptor, respectively. Prior to data collection, single-color controls were used for compensation with Everest software (BioRad). Specifically, three separate samples were used as controls: (1) HEK293T cells expressing mEos3.2, (2) HEK293T cells expressing dsRed2 (as an approximation of a fully photoconverted mEos3.2 sample), and (3) nontransfected HEK293T cells (dark). Data was processed and analyzed using FCSExpress7 software. Data was gated for cells using FSC 488/10-A and SSC 488/10-A, for single cells using FSC 488/10-A and FSC 488/10-H, and for mEos3-expressing cells using 460/22-405nm-A and Donor-A. Finally, the DAmFRET data was plotted in density plots using Acceptor-A on the x-axis and FRET-A/Acceptor-A on the y-axis.

#### 4.2 Determining thresholds to describe cellular states.

DAmFRET provides a powerful tool to visualise the aggregation state of a large population of cells. As discussed in the main text, we expect cells to be in a high or low aggregation state and we observe these two populations in the DAmFRET data. However, since we wish to quantify the fraction of cells in the aggregated state, we need to determine a way to assign each cell in the assay to a state. We do this by introducing a threshold in the scaled fraction of protein in aggregates.

In a DAmFRET dataset, we determine the scaled fraction of protein in an aggregated state, by the ratio of the FRET signal, $F$ divided by the acceptor fluorescence, $A$. Since we are not able to directly convert signals to concentrations - we can only measure the *scaled* fraction. This scaling will be constant within an experiment, but will likely vary across different proteins and cell types.

The scaled protein fraction, which was originally defined as "AmFRET" [9] but which here we call $S$, is given by $S = F/A$. We expect some error on the measurement of both $F$ and $A$ and we assume this is a Gaussian distribution with some constant standard deviation $\sigma_F$ and $\sigma_A$ respectively. For a measured $A$, the spread in $S$ is therefore $\sigma_S = \sigma_F/A$. The 'healthy' state has a low fraction of aggregated protein. We want to determine a threshold in $S$, called $S_T$, that separates most of this low aggregate population from the aggregated state. This ratio of noisy measurements results in the funnel shape seen in the healthy population at low concentrations in the data in the main text Fig. 3d and 5c-e.

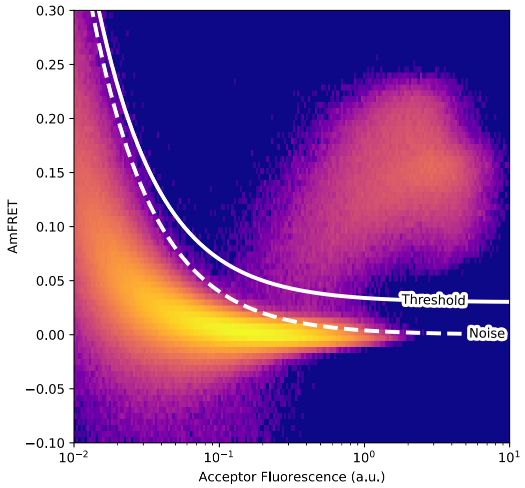


**Figure S7**: Model for noise and threshold boundary between the two states for HET-s (218-289)HET-s (218-289) expressed in yeast. The plot is made by combining the data points for all experiments across all time points (incubation for 16hrs, 24hrs, 32hrs, 48hrs).

###### 4.2.1 Distinct aggregate populations (main text Fig. 3)

For the data in main text Fig. 3, we expect an extremely low aggregate population in the healthy state and any measured signal in this state is noise. We define a threshold at $C + 4\sigma_F/A$ as the boundary between the two states, where $C$ is a constant offset. We can determine $\sigma_F$ by considering the standard deviation of the FRET signal (normalised by SSC) at low protein expression, as we only see the low aggregate population in this region. We consider $0.005 < A < 0.02$ which gives $\sigma_F \approx 1e-3$ using data points across all experiments. Inspecting the DAmFRET data, we set the constant offset as $C = 0.03$ by considering the separation of the two states at large protein expression. Thus we use the threshold $S_T = 0.03 + 0.004/A$ to separate the two regions and, as shown in FigS7 this process well-describes the boundary of the noise and separates the healthy and pathological cases for the experiments used.

###### 4.2.2 Response to preformed fibrils (main text Fig. 4)

We wish to measure via DAmFRET the dose response from treating a population of cells with preformed fibrils. We compare different populations of cells that are all incubated for 48hrs. We

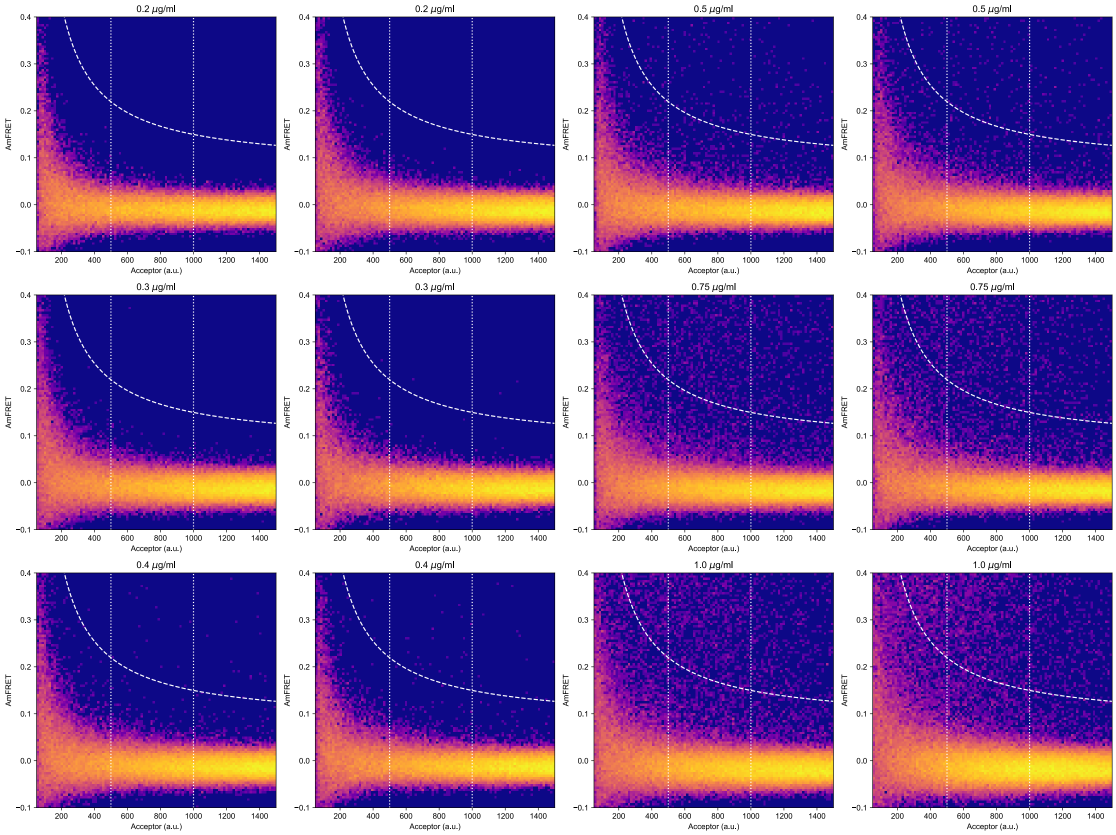


**Figure S8**: Dose dependent response of a cellular population. The dotted lines show the boundaries of region being considered and the dashed line is the threshold for a cell to be considered pathological. All incubations are for 48hrs. Experiments were performed on two different populations for each dose.

choose a specific acceptor fluorescence range to compare between the experiments, which is between 500-1000 a.u. (Fig. S8). As before, we can determine the measurement noise by considering the raw FRET signal at low acceptor fluorescence (< 300 a.u.). In this region we find $\sigma_F \approx 23$ and we set a threshold of $3\sigma_F + C$ where $C$ is a constant offset determined empirically as 0.05.

##### 4.2.3 Response to designed aggregation inhibitors (main text Fig. 4)

We now discuss the thresholding process for cell populations also expressing inhibitory proteins. Again we wish to distinguish a high aggregation and low aggregation state, based on the FRET signal. We include the same affects as described above, we use a threshold at $4\sigma_F$ ($\sigma_F = 3.6$ measured at low acceptor fluorescence for $0.5 < A < 9$ and without normalisation from SSC) and from inspection we set $C = 0.08$. However, at high protein expression, monomeric A$\beta$ begin to oligomerise into non-amyloid structures and thus expected to transiently be close together and thus generate a signal without being part of an aggregate, which we call non-amyloid FRET. The contribution of this effect will scale as $A^2$, consistent with what is seen in the data. We illustrate this by combining all data sets and showing a log-log plot of the fluorescence distribution, see Fig. S9a. This shows two separate populations with the slope of the population maximum intensities $\approx 2$, consistent with the reaction order expected for the non-aggregate FRET effect. We can align the threshold with the separation between the two regions in this regime and use the threshold $S_T = 0.08 + (13.6 + 5 \cdot 10^{-5} A^2)/A$. Combining all data binder sets, as shown in Fig. S9, this works well at both high (visible in (a)) and low (visible in (b)) acceptor fluorescence. We now discuss the thresholding process for cell populations also expressing inhibitory peptides. Again we wish to distinguish a high aggregation and low aggregation state, based on the FRET signal. We include the same affects as described above, we use a threshold at $4\sigma_F$ ($\sigma_F = 3.6$ measured at low acceptor fluorescence for $0.5 < A < 9$ and without normalisation from SSC) and from inspection we set $C = 0.08$. However, at high protein expression, monomeric A$\beta$ may begin to transiently interact in the form of non-amyloid structures. This may generate a weak FRET signal without runaway aggregation, which we call non-amyloid FRET. The contribution of this effect will scale as $A^2$, consistent with what is seen in the data. We illustrate this by combining all data sets and showing a log-log plot of the fluorescence distribution, see Fig. S9a. This shows two separate populations with the slope of the population maximum intensities $\approx 2$, consistent with the reaction order expected for the non-aggregate FRET effect. We can align the threshold with the separation between the two regions in this regime and use the threshold $S_T = 0.08 + (13.6 + 5 \cdot 10^{-5} A^2)/A$. Combining all data binder sets, as shown in Fig. S9, this works well at both high (visible in (a)) and low (visible in (b)) acceptor fluorescence.

We wish to understand how the binders change the protein expression at which the cell population changes from mostly unaggregated protein to mostly aggregated protein. We can bin the cells within a specific acceptor fluorescence and consider the fraction of cells above the threshold within each bin. The critical tipping point would suggest this fraction changes from 0 to 1 at some specific protein expression, however due to cell-to-cell variability we might expect this transition to be more sigmoidal. Additionally, for the A$\beta$ data in main text Fig. 5, the maximum fraction of aggregated

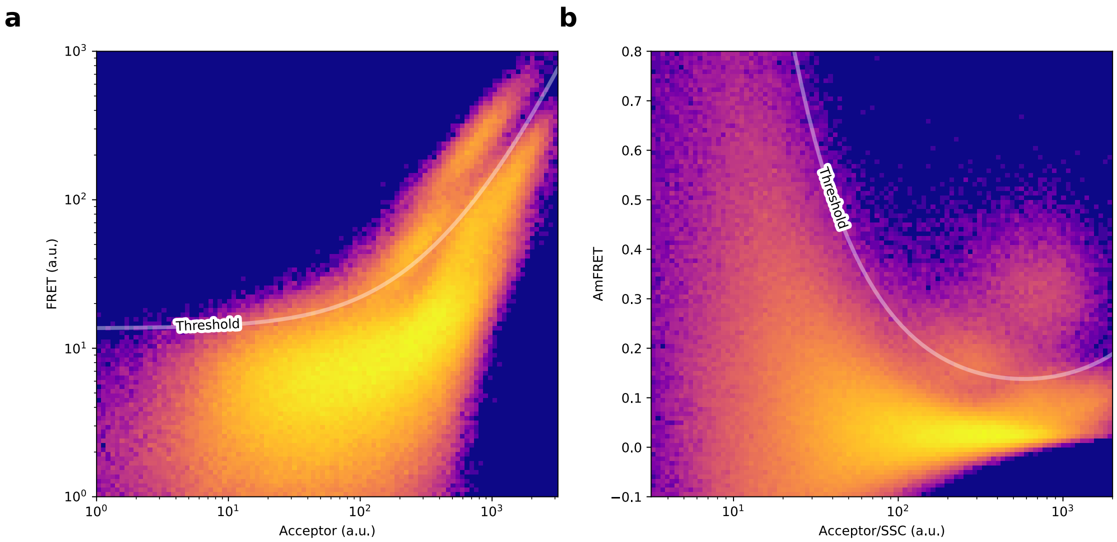


**Figure S9**: Complete noise thresholding model including non-aggregate FRET. The data shown is from combining the three systems of Weak, Strong and No Inhibitors across all repeats (N=3). The raw FRET signal, (a), has not been normalised by the total protein expression.

cells plateaus below 1 even at large fluorescence values. We believe this is partly a measurement artifact: we measure here the total fluorescence rather than the concentration of protein (as we presently lack a suitable proxy for HEK cell volume), thus large cells with a lower concentration will appear at the same x position as smaller cells with a higher protein concentration. This may lead to some cells at high fluorescence values actually being below the critical monomer concentration, which in turn results in a plateau in the fraction aggregated that is below 1. We expect that the transition point, however, still provides a good estimate for the critical monomer concentration. To extract its value, we renormalise the data of the fraction of aggregated cells (parameters in Table S7) and fit the equation $y = 1/(1 + \exp(q(A_{1/2} - A))$, where $y$ is the renormalised fraction of aggregated cells, using the software AmyloFit (*26*) to fit $q$ and $A_{1/2}$. The fitting parameters are given in Table S7.

#### 4.3 Fitting kinetic rates in the presence of inhibitors.

The experimental details of the kinetic assays and purification of the binders are given in Sahtoe et al. (*47*). Briefly, 2 $\mu$M monomeric A$\beta$42 are aggregated in the presence of different concentrations of binder, the increase in aggregate mass over time is tracked using Thioflavin T fluorescence. From these data, the effective rates of aggregation in the presence and absence of inhibitor were determined from the aggregation kinetics as detailed in Meisl et al. (*26*). We here use the effective rates obtained in the presence of 1 $\mu$M binder, as these most closely resemble the situation in the cell assay (inhibitor is expressed at concentrations comparable to, but slightly lower than the monomer).

#### 4.4 Relating critical concentration to effective rates.

To obtain an approximate expression for the critical monomer concentration, $m_{\text{crit}}$, in our cellular assay, we approximate the removal and production terms in equation (1) of the main text as $A(\mu, m_{\text{tot}}) = \kappa M$ and $R(\mu, m_{\text{tot}}) = \lambda M$, where $M$ is the aggregate mass and $\kappa$ is the monomer concentration dependent rate of aggregation as defined in previous work (*23, 26*). This is equivalent to the assumption that production is dominated by the self-replication process, a reasonable assumption for A$\beta$, and that the removal rate is proportional to aggregate mass. The critical point is then given by $\lambda = \kappa$ (*16*). To determine the monomer concentration at which this transition occurs, we need to explicitly account for the monomer dependence of $\kappa$. We assume that the aggregate removal rate $\lambda$ is independent of the monomer concentration. Experimentally, the concentration dependence of $\kappa$, i.e. the reaction order of aggregation with respect to the monomer concentration, ranges from 0.5 to 1.5 for A$\beta$42, depending on the degree of saturation of secondary nucleation (*48, 49, 67*). As a simple approximation, we can thus write $\lambda = k_{\text{eff}} m_{\text{crit}}^{\gamma}$, where $\gamma$ is the reaction order and $k_{\text{eff}}$ an effective aggregation rate, taking into account inhibition. Rearranging and taking logarithms gives $\log(\frac{\lambda}{k_{\text{eff}}}) = \gamma \log(m_{\text{crit}})$.

If this effective aggregation rate in cells is proportional to the effective aggregation rate in vitro, we thus expect a double logarithmic plot of the effective rate from in vitro experiments, $k_{\text{eff}}$, against the critical monomer concentration from cell experiments, $m_{\text{crit}}$, to fall on a straight line, with slope given by $\gamma$, as demonstrated in the main text.

### 5 Effect of the Introduction of Preformed Seeds

#### 5.1 Multi-hits in a Poisson Process

In a nucleation-limited cell, a single aggregate can trigger the transition from an unaggregated to aggregated state. In a system where removal is balanced with a low aggregation rate, it is possible that multiple aggregates are required to trigger the transition to runaway aggregation. We assume that the arrival and entry of seed aggregates into cells is independent and occurs at a constant rate. In line with established kinetics, this rate will be proportional to the concentration of seed in solution that is dosed onto cells.

The arrival/entry times of a seed are therefore governed by Poisson statistics so that the probability of $x$ seeds arriving during an experiment is

$$p(x \text{ seeds arrive during experiment}) = \frac{(vC)^x e^{-vC}}{x!} \tag{S17}$$

where $C$ is the seed concentration and $v$ is the entry/arrival rate constant. In a system were uptake is less efficient, $v$ will be lower, however the total cellular uptake rate will remain proportional to $C$.

The multi-hit describes the probability of $h$ or more events during an experiment, which equation (S17) gives as

$$p(h \text{ or more events}) = 1 - p(\text{fewer than } h \text{ events}) = 1 - e^{-vC}\sum_{x=0}^{h-1}\frac{(vC)^x}{x!}. \tag{S18}$$

For a fixed $h$, we can fit for $v$ using a dose response curve that shows the fraction of cells that have aggregated, $p$, by treatment withy some dose of seeds, $C$. When the dose is low we expand (S18) in $vC$ to get

$$p(n \text{ or more events}) \approx \frac{(vC)^h}{h!} + O((vC)^{h+1}). \tag{S19}$$

Thus, at small doses, we find that the fraction of aggregated cells $\propto C^h$. This can be easily read of a log-log plot and the equation (S18) is plotted in Fig. S10. We see that the different $h$ hits can be easily identified from the slope of the line, panel (a), and that a different rate of uptake or fraction of competent seeds (which only affect $v$) only act to translate the line, but they don't change the slope.

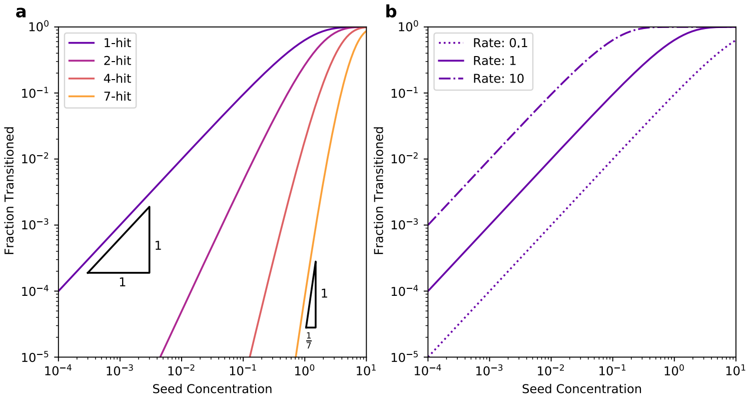


**Figure S10**: Slope variation for different *n*-hit behaviour.

#### 5.2 Fitting of Seeding Data

We fit (S18) to the data in Fig. 2 of the main text. We use a Nelder-Mead simplex algorithm to minimise the sum of the square log-space $y$ differences between the functional form and the measured data. We do this for different discrete hit numbers (typically from 1 to 9) and determine the best global fit. We report the parameters for the best fit as well as references to the experimental data in Table S8.

**Table S1**: Frequently used parameter values for determining stability lines.

| Variable | Value |
|---|---|
| $k_n$ | $1.0 \times 10^3 \mathrm{M}^{1-n_c}\mathrm{hr}^{-1}$ |
| $k_+$ | $1.2 \times 10^8 \mathrm{M}^{-1}\mathrm{hr}^{-1}$ |
| $n_c$ | 2.0 |
| $n_2$ | 2.0 |
| $k_2$ | $1.2 \times 10^{14} \mathrm{M}^{-n_2}\mathrm{hr}^{-1}$ |
| $\lambda$ | $1 \times 10^4 \mathrm{hr}^{-1}$ |
| $K_\lambda$ | $1 \times 10^{-10} \mathrm{M}$ |

**Table S2**: Fitting of healthy control parameters. Lower and upper are 68th percentile confidence intervals. Samples refer to number of measured aggregates.

| Protein | Fitting parameters | | $\log(\delta_H)$ | | | Samples |
|---|---|---|---|---|---|---|
| | $i_{min}$ | $i_{max}$ | Lower | Mean | Upper | |
| Tau | 500 | 2000 | -2.48 | -2.47 | -2.46 | 1684 |
| $\alpha$-synuclein | 500 | 1500 | -2.24 | -2.22 | -2.21 | 1755 |

**Table S3**: Fitting of disease parameters. Lower and upper are 68th percentile confidence intervals. Samples refer to number of measured aggregates.

| Condition | Protein | Fitting parameters | | $r_C$ | | | $\beta$ | | | Samples |
|---|---|---|---|---|---|---|---|---|---|---|
| | | $i_{min}$ | $i_{max}$ | Lower | Mean | Upper | Lower | Mean | Upper | |
| AD | Tau | 500 | 4000 | 0.36 | 0.38 | 0.38 | 0.86 | 0.89 | 0.91 | 13720 |
| CBD | Tau | 500 | 1750 | 0.62 | 0.78 | 0.93 | 0.09 | 0.38 | 0.71 | 2241 |
| PSP | Tau | 500 | 2000 | 0.68 | 0.75 | 0.81 | 0.53 | 0.73 | 0.92 | 3738 |
| Picks | Tau | 500 | 2000 | 0.71 | 0.75 | 0.78 | 0.75 | 0.87 | 0.97 | 5077 |
| PD | $\alpha$-synuclein | 500 | 3000 | 0.36 | 0.39 | 0.40 | 0.78 | 0.82 | 0.85 | 5225 |

**Table S4**: Tauopathy Patient Information for the length distribution data.

| Diagnosis | Case Number | Sex | Age at death (years) | PMI (h) | Region | Case ID |
|---|---|---|---|---|---|---|
| AD | 1 | F | 72 | 66 | BA6/8 | BBN001.36924 |
| AD | 2 | M | 75 | 71 | BA6/8 | BBN001.37400 |
| AD | 3 | F | 85 | 45 | BA6/8 | BBN_25739 |
| AD | 4 | M | 75 | 66 | BA6/8 | BBN001.36839 |
| AD | 5 | M | 75 | 92 | BA6/8 | BBN001.36689 |
| PSP | 1 | M | 73 | 103 | BA46 | BBN001.36927 |
| PSP | 2 | M | 80 | 101 | BA46 | BBN001.37213 |
| PSP | 3 | M | 81 | 78 | BA46 | BBN001.37236 |
| PSP | 4 | M | 70 | 66 | BA46 | BBN001.35381 |
| PSP | 5 | M | 70 | 46 | BA46 | BBN001.37087 |
| PiD | 1 | M | 71 | 58 | BA9/10/46 | N/A |
| PiD | 2 | M | 72 | 37 | BA9/10/46 | N/A |
| PiD | 3 | M | 68 | 67 | BA9/10/46 | N/A |
| PiD | 4 | M | 74 | 102 | BA9/10/46 | N/A |
| CBD | 1 | F | 73 | 67 | BA9/10/46 | N/A |
| CBD | 2 | M | 75 | 74 | BA9/10/46 | N/A |
| CBD | 3 | F | 76 | 51 | BA9/10/46 | N/A |
| CBD | 4 | F | 79 | 88 | BA9/10/46 | N/A |
| CBD | 5 | F | 82 | 78 | BA9/10/46 | N/A |
| HC | 1 | F | 71 | 95 | BA6/8 | BBN001.29882 |
| HC | 2 | M | 72 | 60 | BA6/8 | BBN001.30178 |
| HC | 3 | F | 73 | 74 | BA6/8 | BBN001.35138 |
| HC | 4 | M | 71 | 71 | BA6/8 | BBN001.30916 |
| HC | 5 | M | 82 | 56 | BA6/8 | BBN001.35549s |

**Table S5**: Patient Information for the $\alpha$-synuclein length distribution data.

| Case | Condition | Stage | Age at death (years) | Sex | Region |
|---|---|---|---|---|---|
| 1 | Control | 0 | 70 | Male | BA10-11 |
| 2 | Control | 0 | 70 | Female | BA10-11 |
| 3 | Control | 0 | 82 | Female | BA10-11 |
| 4 | PD | 4 | 86 | Female | BA10-11 |
| 5 | PD | 4 | 78 | Female | BA10-11 |
| 6 | PD | 5 | 74 | Male | BA10-11 |

**Table S6**: Reagent Table for DAmFRET experiments.

| Reagent/Resource | Source | Catalog number |
|---|---|---|
| DMEM, high glucose, pyruvate | ThermoFisher | 11995073 |
| FUGENE HD | Promega | E2312 |
| Opti-MEM™I Reduced Serum Medium | ThermoFisher | 31985062 |
| Premium Grade Fetal Bovine Serum (FBS), Heat Inactivated | VWR | 97068-091 |
| Gibco™ Penicillin-Streptomycin | FisherScientific | 15-140-122 |
| Phosphate-Buffered Saline (PBS) without calcium and magnesium | Corning | 21-040-CV |
| TrypLE | Gibco | 12604-021 |
| Lipofectamine™ 3000 Transfection Reagent | Thermofisher | L3000015 |
| 4% Paraformaldehyde (PFA) in 1X PBS | ThermoFisher | J19943-K2 |
| 96-well round-bottom plate | Corning | 3359 |
| 12-well plate | Corning | 3513 |

**Table S7**: Result of fitting the binder aggregation profiles. The data was fit using all repeats (N=3). parameters from binder fitting parameters.

| Data Set | Normalisation Offset | Normalisation Constant | q (a.u.$^{-1}$) | $A_{1/2}$ (a.u.) |
|---|---|---|---|---|
| No Inhibitor | 0.010 | 0.36 | 0.023 | 190 |
| Weak Inhibitor | 0.0091 | 0.44 | 0.0052 | 660 |

**Table S8**: Best fit parameters and dataset references for seeding data in Fig. 4 of the main text.

| Dataset Name | Hit Number | Rate | Reference |
|---|---|---|---|
| Virus | 1 | $3.52 \times 10^{-12}$ | (*37*) |
| OHSC1 | 6 | $3.72 \times 10^{-2}$ | (*38*) |
| OHSC2 | 7 | $3.47 \times 10^{-2}$ | (*37*) |
| M$\beta$CD | 3 | $8.26 \times 10^{-2}$ | (*38*) |
| Tau DAmFRET | 5 | 3.31 | See section 4 |
